# Changes of Mutations and Copy-number and Enhanced Cell Migration during Breast Tumorigenesis

**DOI:** 10.1101/2022.03.27.485986

**Authors:** Seung Hyuk T. Lee, Joon Yup Kim, Peter Kim, Zhipeng Dong, Chia-Yi Su, Eun Hyun Ahn

## Abstract

Although cancer stem cells (CSCs) play a major role in tumorigenesis and metastasis, the role of genetic alterations in invasiveness of CSCs is still unclear. Tumor microenvironment signals, such as extracellular matrix (ECM) composition, significantly influence cell behaviors. Unfortunately, these signals are often lost in *in vitro* cell culture. This study determines putative CSC populations, examines genetic changes during tumorigenesis of human breast epithelial stem cells, and investigates single-cell migration properties on ECM-mimetic platforms. Whole exome sequencing data indicate that tumorigenic cells have a higher somatic mutation burden than non-tumorigenic cells, and that mutations exclusive to tumorigenic cells exhibit higher predictive deleterious scores. Tumorigenic cells exhibit distinct somatic copy number variations (CNVs) including gain of duplications in chromosomes 5 and 8. ECM-mimetic topography selectively enhances migration speed of tumorigenic cells, but not of non-tumorigenic cells, and results in a wide distribution of tumorigenic single-cell migration speeds, suggesting heterogeneity in cellular sensing of contact guidance cues. This study identifies mutations and CNVs acquired during breast tumorigenesis, which can be associated with enhanced migration of breast tumorigenic cells, and demonstrates that a nanotopographically-defined platform can be applied to recapitulate an ECM structure for investigating cellular migration in the simulated tumor microenvironment.

## 1. Introduction

Cancer stem cells (CSCs) account for rapid recurrence and therapeutic resistance after primary cancer treatment, posing difficulties in treatment strategies^[1,2]^. CSCs contribute to tumor heterogeneity and metastatic dissemination^[1]^. Breast CSCs have been demonstrated to mediate cancer cell invasiveness and metastasis^[1,3]^. A subpopulation of breast CSCs characterized with CD44^+^/CD24^-/low^ are more resistant to therapy and are more invasive than breast non-CSCs^[4#x2013;6]^. Contact guidance of extracellular matrix (ECM) anisotropy that recapitulates a tumor microenvironment is an important factor in modeling tumor progression and metastasis^[7#x2013;9]^. Over the past several years, advances in the fabrication of ECM-mimetic nanoscale topography have facilitated the study of how cells sense contact guidance cues and of cellmatrix interactions^[8,10–13]^. However, little is known about how the contact guidance of cell migration is affected at a single-cell level by tumorigenicity.

Genetic alterations in CSCs, such as somatic point mutations and copy number variations (CNVs), have been associated with cancer cell migration and heterogeneity^[14]^. Somatic mutations in different regions of the genome can lead to distinct subtypes and subpopulations of cancer cells^[15]^ and may impact how cells sense and respond to contact guidance cues differently^[11]^. Despite this importance, studies investigating cancer cell migration and metastasis using ECM-mimetic platforms often lack genomic profiles of the cells investigated. Examining genomic differences between cells representing different stages of tumorigenesis and subpopulations, particularly CSC, together with ECM-mimetic platforms could help to identify potential genetic mechanisms underlying cancer cell migration and invasiveness.

To study the effect of anisotropic contact guidance on cell migration properties, we investigated a human breast epithelial cell (HBEC) model. Our model cells represent multi-stages of breast carcinogenesis, which were sequentially derived from the same parental normal breast epithelial cells with stem cell features. Development, characterization, and culture of these transformed HBECs has been described^[16#x2013;27]^ and is highlighted in the *2.1 section* of *Materials and Methods.* In this study, immortalized non-tumorigenic cells (referred to as ‘non-tumorigenic cells’ hereinafter) and highly tumorigenic-xenograft cells (‘tumorigenic cells’) were examined.

Our flow cytometry results indicate that the tumorigenic cells have a 10-fold higher putative CSC population than did non-tumorigenic cells. We examined single base substitutions (point mutations) and CNVs in tumorigenic and non-tumorigenic cells using whole-exome sequencing (WES). WES revealed a higher somatic mutation burden in tumorigenic cells than non-tumorigenic cells and identified somatic mutations in genes associated with breast cancer cell proliferation and metastasis. In addition, we observed significant differences in CNVs between non-tumorigenic and tumorigenic cells, and these differences are more frequently observed in duplications than in deletions. Using a nanotopographically-defined ECM-mimetic platform, we found that anisotropic contact guidance is required to trigger the highly migratory phenotype of tumorigenic cells. The broad distribution of migration speed in tumorigenic cells suggests heterogeneity in cellular sensing of contact guidance cues. Our findings highlight the critical roles of tumor ECM topography in the migration of tumorigenic stem cell-like cells.

## 2. Materials and Methods

### 2.1. Human breast epithelial cells (HBECs) and *in vitro* transformed HBECs

The development and culture of normal HBECs have been described previously ^[18,22,23]^. Immortalized non-tumorigenic, weakly tumorigenic, highly tumorigenic, and highly tumorigenic xenograft cells were transformed sequentially from the same parental normal stem cells with SV40 large T-antigen, x-rays, and the *ERBB2* oncogene^[20#x2013;22]^. Highly tumorigenic cells were injected into nude mice and then the tumors formed in nude mice were collected and grown in culture to develop highly tumorigenic xenograft (M13SV1R2N1-Xeno) cells.

Development, characterization, and culture of these transformed HBECs have been described previously^[16#x2013;27]^. Immortalized non-tumorigenic (M13SV1) cells and highly tumorigenic-xenograft (M13SV1R2N1-xeno) cells were examined in this study. Mutations in normal HBECs with stem cell features (HME13) (previously referred as Type I HBECs)^[18,22]^, that we identified using WES, were used to filter out germline mutations in the non-tumorigenic and tumorigenic cells. The cells (M13SV1R2N1-Xeno, M13SV1, and HME13) were provided by Dr. Chia-Cheng Chang at Michigan State University (MSU) in East Lansing, MI, USA. A material transfer agreement (MTA) was approved by both MSU and the University of Washington (UW) and experiments were performed in UW. The cells were authenticated by short tandem repeat (STR) DNA profiling (Genetica DNA Laboratories, Labcorp brand, Burlington, NC, USA).

### 2.2. Identification of breast cancer stem-like cell population using flow cytometry

Cells were collected two days after initial seeding and were incubated with antibodies labeled with fluorochromes anti-CD24-PE and anti-CD44-APC (BD Biosciences, USA). Cell sorting and immunofluorescence analysis were performed using BD FACS Aria or BD FACSCanto II (BD Immunocytometry Systems). Cell debris and doublets were excluded by using forward and side scatter functions of the FACS instrument. The viable breast cancer stem cell (CSC) population (CD44+/CD24-/low)^[16,24,28]^ was calculated using the FlowJo program version 9.5 (Tree Star, Inc).

### 2.3. DNA library preparation and whole-exome sequencing (WES) data processing and analysis

WES procedures for DNA extraction, adapter synthesis, library preparation, and exome capture were carried out as previously described^[29,30]^. DNA was lysed with a lysis buffer (10 mM Tris-HCl, pH 8.0, 150 mM NaCl, 20 mM EDTA, 1% SDS) and proteinase K (Qiagen) was added to samples to degrade proteins. Then DNA was extracted using UltraPure Phenol:Chloroform:Isoamyl Alcohol (Invitrogen, Thermo Fisher Scientific Inc., Carlsbad, CA, USA). The DNA libraries of normal HBECs (HME13), non-tumorigenic cells (M13SV1), and tumorigenic-xenograft cells (M13SV1R2N1-xeno) were captured using SeqCap EZ Exome v2 (Nimblegen) and hybridized to biotinylated capture probes for paired-end sequencing on an Illumina HiSeq 2500 platform (Illumina Inc.).

Variants were called using the human reference genome hg19 from the University of California Santa Cruz, and only chromosomal positions cited in our exome target BED file were kept. In addition, base positions with unreadable counts >10% of total reads or read coverage below 20 were excluded. Sequence reads were aligned to the targeted exome using BWA v0.6.1-r104 and then were realigned using GATK v1.5-21. The aligned reads were controlled for accuracy by filtering out mapped reads with a MAPping Quality (MAPQ) value below 20 (genome.sph.umich.edu/wiki/Mapping_Quality_Scores). Bases with low calling accuracy (Phred quality score below 13, meaning an error probability of 5%) (www.illumina.com/science/technology/next-generation-sequencing/plan-experiments/quality-scores.html) were not considered for variant calling.

To exclude batch specific sequence artifacts, only positions with a variant read of 5 or greater in at least one sample out of all samples sequenced and prepared from the same WES DNA library preparation experiment were included (i.e., only positions with maximum variant read of 5 or greater among all samples sequenced in the same batch were included). This procedure is analogous to retaining variants representing at least 5% of total reads given average coverage of 100 across all samples sequenced in the same batch. In addition, germline mutations were removed in non-tumorigenic and tumorigenic cells by removing mutations found in the matching normal breast epithelial stem cells.

Mutations in non-tumorigenic and tumorigenic cells were annotated with predicted deleterious scores by CHASMplus, CHASMplus-BRCA^[31]^, and CADD Exome^[32]^ using OpenCRAVAT^[33]^. Pathogenicity scores by CHASMplus and CHASMplus-BRCA range between 0 and 1, with higher scores indicating a greater likelihood of being a driver mutation. CHASMplus scores were derived using the 32 cancer types from The Cancer Genome Atlas (TCGA), and CHASMplus_BRCA scores were trained using Breast Invasive Carcinoma samples from TCGA.

Somatic copy number variations (CNVs) in non-tumorigenic and tumorigenic cells were determined using CopyDetective^[34]^, an algorithm that performs threshold (cell fraction) aware CNV calling for WES data in each sample against a matched control. The same parental normal stem cells (HME13) of non-tumorigenic and tumorigenic cells were used as the matching control to call CNVs in autosomes. The CNVs were represented as a percent of cell fraction in non-tumorigenic or tumorigenic cells and were assigned with quality scores calculated from the single nucleotide polymorphisms (SNPs) affected by the CNVs. All SNPs present in any of normal stem, non-tumorigenic, and tumorigenic cells at base positions with minimum combined read coverage of 20 were included. Raw CNV calls with cell fractions less than ~5%, or with quality scores less than the default threshold were filtered out. Then these filtered CNV calls were merged based on their genomic positions and further filtered with a default quality score threshold of 10.76 to identify regions of significant changes of CNVs in comparison to the matched control.

### 2.4. Fabrication of cell adhesion substrate mimicking nanometer-scale features of the extracellular matrix (ECM)

Cells were cultured on surfaces made from the UV-curable, biocompatible polymer polyurethane-acrylate-301 (PUA) (Minutatek, South Korea). Nanopatterns were generated on a polyethylene terephthalate (PET) film using a silicon master template and UV-assisted capillary force lithography (CFLA)^[35]^. The nanopatterned PET was used as a template to create nano-grooves on a primer-treated cover-glass using PUA and UV-assisted CFLA. Each nano-groove (nanopatterned) measured 800nm wide and 800nm high was separated from its neighbor by 800nm to mimic the ~850nm mean spacing between ECM collagen fibril bundles^[8,11]^. The unpatterned surface was coated with a layer of PUA and cured with UV prior to plasma and collagen treatment to produce a non-textured surface coat. Both unpatterned (UP) and nanopatterned (NP) surfaces were made hydrophilic via treatment with plasma oxygen for 5 minutes at 100W power and with 100 sccm of O2 at 0.5 torr using a Femto Science SUTE-MPR Plasma treatment machine. Then, to further enhance cellular attachment^[36]^, the plasma-treated surfaces were coated with 35 μg/mL of Collagen I Rat Protein, Tail (cat# A1048301, Gibco Thermo Fisher Scientific, Waltham, MA, USA) and incubated at 37°C for 3 hours.

### 2.5. Live cell imaging

Non-tumorigenic and tumorigenic human breast epithelial cells were seeded on unpatterned and nanopatterned PUA301 surfaces for six hours (17 hours for the supplementary data experiment) and were then imaged for 12 hours using a Nikon Eclipse Ti microscope. Images of the cells were taken for 12 hours at 24 spots for each group, for a total of 96 spots, at the intervals of 20 minutes for the main experiment (or 10 minutes for the supplementary data experiment). The cells were incubated at 37°C and 5% CO2 during the live-cell imaging. The images taken at different time points were sequentially compiled to generate a movie file for each spot imaged.

### 2.6. Cell migration quantification

Individual cells were tracked to quantify their speed, persistence, and trajectory using the ImageJ program, version 1.49 (https://imagej.nih.gov/ij/). For each movie of migrating cells, the Manual Tracking plugin was used to establish coordinate points, distance, velocity, and pixel values of individual cells at each time point. These established data were then used with in-house MATLAB scripts to calculate speed and persistence and to generate trajectory graphs of each manually tracked single-cell, as described previously^[11]^. Briefly, we calculated speed of each migrating cell by dividing the root mean squared displacement (MSD) of each migrating cell based on N sequential positions tracked by a constant time interval Δt. Total of N = 36 sequential positions were tracked at a time interval Δt = 20 minutes for the main experiment (or N = 36 and Δt = 10 minutes for the supplementary data experiment). Migration speeds of the same cell type seeded on the same surface (unpatterned *vs.* nanopatterned) were averaged. Only single, viable cells that did not divide and that survived during the entire time of live cell imaging were included for the single-cell migration quantification.

### 2.7. Statistical analysis

Differences in the average CHASMplus scores for mutations exclusive to non-tumorigenic (n=95) or tumorigenic cells (n=65) available with a CHASMplus score were analyzed by a two-sided Mann-Whitney U-test from SciPy 1.7.1^[37]^. Differences in the average migration speed between the two groups were analyzed by the ANOVA on Ranks test using SigmaPlot (version 12.0, Systat Software, Inc., San Jose, CA, USA). Differences in the distributions of individual cell migration speed were analyzed by a twosided Kolmogorov-Smirnov test using R-3.6.3. Differences between the groups for all statistical tests were considered significant at *p* < 0.05. Sample size for each ANOVA on Ranks test and the Kolmogorov-Smirnov test can be found in the corresponding figure legends.

## 3. Results

### 3.1. Higher percentages of putative breast CSCs are present in tumorigenic cells than in non-tumorigenic cells

Putative breast CSC populations were determined using the well-known breast CSC marker CD44^+^/CD24^-/low[16,26]^ via flow cytometry (**Figure 1**). These data indicate that tumorigenic cells contain a higher percentage of CSCs (~27.3%) than in non-tumorigenic cells (~2.78) (**Figure 1**) by about 10-fold.

**Figure 1.**
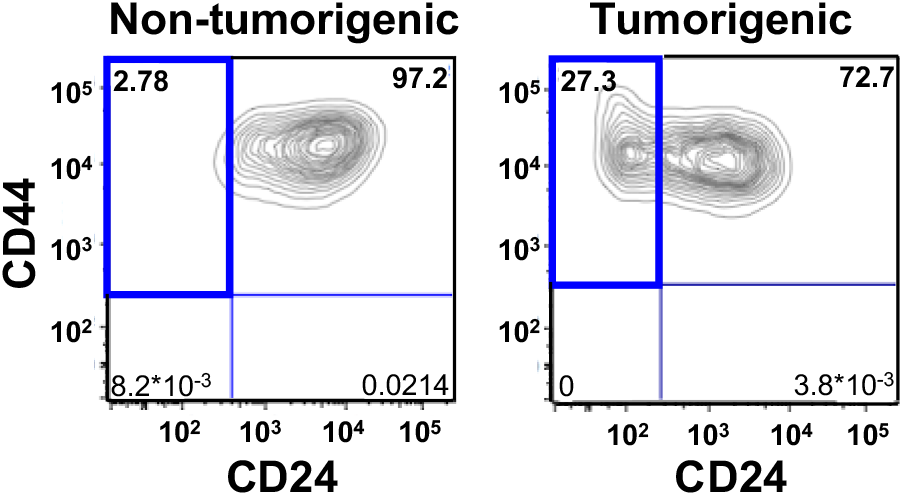
The percentages of cells in each of four subpopulations, defined by the expressions of CD44 and CD24, were determined using flow cytometry in human breast non-tumorigenic and tumorigenic cells. Bold and blue lines represent the percentage of putative breast CSC populations (CD44^+^/CD24^-/low^).

### 3.2. Tumorigenic cells are more highly mutated in genes associated with breast cancer

WES was performed on the DNA of normal stem, non-tumorigenic and tumorigenic cells. Somatic mutations in non-tumorigenic and tumorigenic cells were identified after filtering out matching variants present in normal stem cells, which are presumably germline mutations. Only mutations with variant allele frequency (VAF) of at least 5% were included. A total of 175 point mutations across 133 genes were present in the non-tumorigenic cells but not in the tumorigenic cells (**Figure 2A, Table S1A**). Nonsynonymous mutations were most common in the non-tumorigenic cells followed by synonymous, untranslated region (UTR), and intronic mutations.

**Figure 2.**
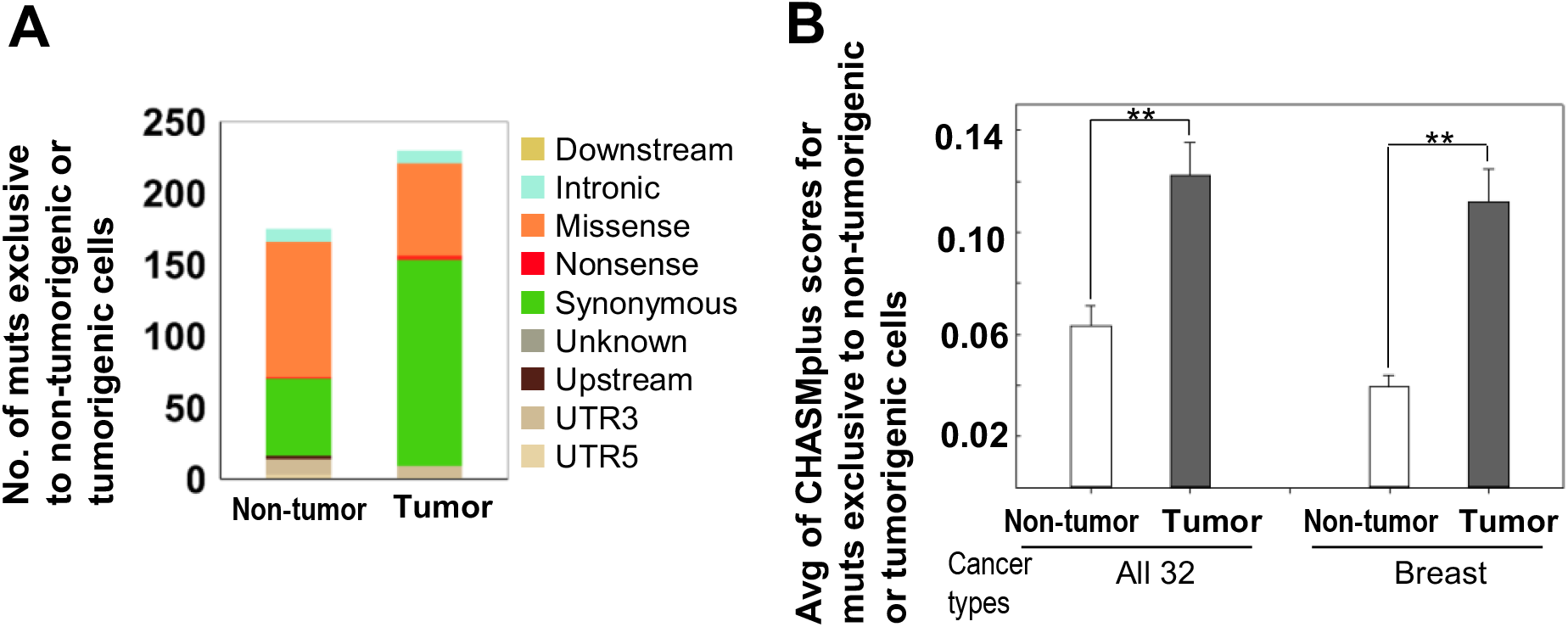
Mutations exclusive to human breast non-tumorigenic or tumorigenic cells were identified using whole exome sequencing. Mutations (muts) with variant allele frequencies less than 5% and mutations found in the matched normal cells (HME13) were filtered out. **(A)** Numbers of mutations exclusive to non-tumorigenic or tumorigenic cells. Each color represents proportion of each category of point mutations annotated using ANNOVAR. Abbreviations used are: UTR3, three prime untranslated region; UTR5, five prime untranslated region. **(B)** Averages (Avg) of CHASMplus scores based on all 32 cancer types and on breast cancer alone for mutations exclusive to non-tumorigenic (n=65) or tumorigenic cells (n=95) available with CHASMplus scores. Errors bars indicate standard error of the mean. Significant differences in the averages of CHASMplus scores are indicated (**: *p*<5×10^-4^) by the Mann– Whitney U-test.

A higher somatic mutation burden was observed in the tumorigenic cells with a total of 230 point mutations across 81 genes (**Figure 2A, Table S1B**). In contrast to the non-tumorigenic cells, synonymous mutations were most common in the tumorigenic cells, followed by nonsynonymous, UTR, and intronic mutations. Most of the mutations present exclusively to the tumorigenic cells were found in the gene *ERBB2*, which is related to a transformation step of the *ERBB2* oncogene overexpression (**Figure S1, Table S1B**) introduced during development of this cell model. Other mutations were identified in genes that have been previously reported to be associated with breast cancer, such as *FAACD2*^[38,39]^, *MMP13*^[40]^ *MSH2*^[41]^, and *RET*^[42]^.

### 3.3. Predicted pathogenicity scores are higher in mutations exclusive to tumorigenic cells than in mutations exclusive to non-tumorigenic cells

From the total of 175 and 230 mutations exclusive to non-tumorigenic or tumorigenic cells, respectively, we annotated 95 non-tumorigenic exclusive mutations and 65 tumorigenic exclusive mutations with CHASMplus scores^[31]^ to predict pathogenicity. The scores were generated based on all 32 cancer types and on breast cancer alone (**Figure 2B**). The number of genes that have mutations annotated with the CHASMplus scores are identical in both settings (32 cancer types; breast cancer alone). The genes with high CHASMplus scores mutations such as *ITCH*^[43]^, *CHD8*^[44]^ and *SREBP-2*^[45]^ have also been linked to breast cancer cell proliferation and metastasis.

### 3.4. Mutations that became increasingly mutated during tumorigenesis of HBEC are identified

Among mutations present in both non-tumorigenic and tumorigenic cells, we identified mutations that became increasingly mutated (clonally expanded) during tumorigenesis of HBECs. A total of 74 mutations were identified on exons of 54 genes with higher VAFs in tumorigenic cells than in non-tumorigenic cells by at least 1.5-fold (**Figure 3, Table S1C**). Most of the identified mutations are C>T/G>A transitions, followed by C>G/G>C transversions and T>C/A>G transitions (**Figure 3A, Table S1C**). Among these 74 mutations, the mutations with the three highest scores of CHASMplus-all-cancers (0.164, 0.129, and 0.124) and of CHASMplus-breast-cancer (0.068, 0.073, and 0.034) are: C>T in *CHD1*, C>A in *GNAS,* and A>G in *CYBA,* respectively (**Figure 3A, Table S1C**). These top three mutations are missense mutations (E986K, A436D, and V174A, respectively).

**Figure 3.**
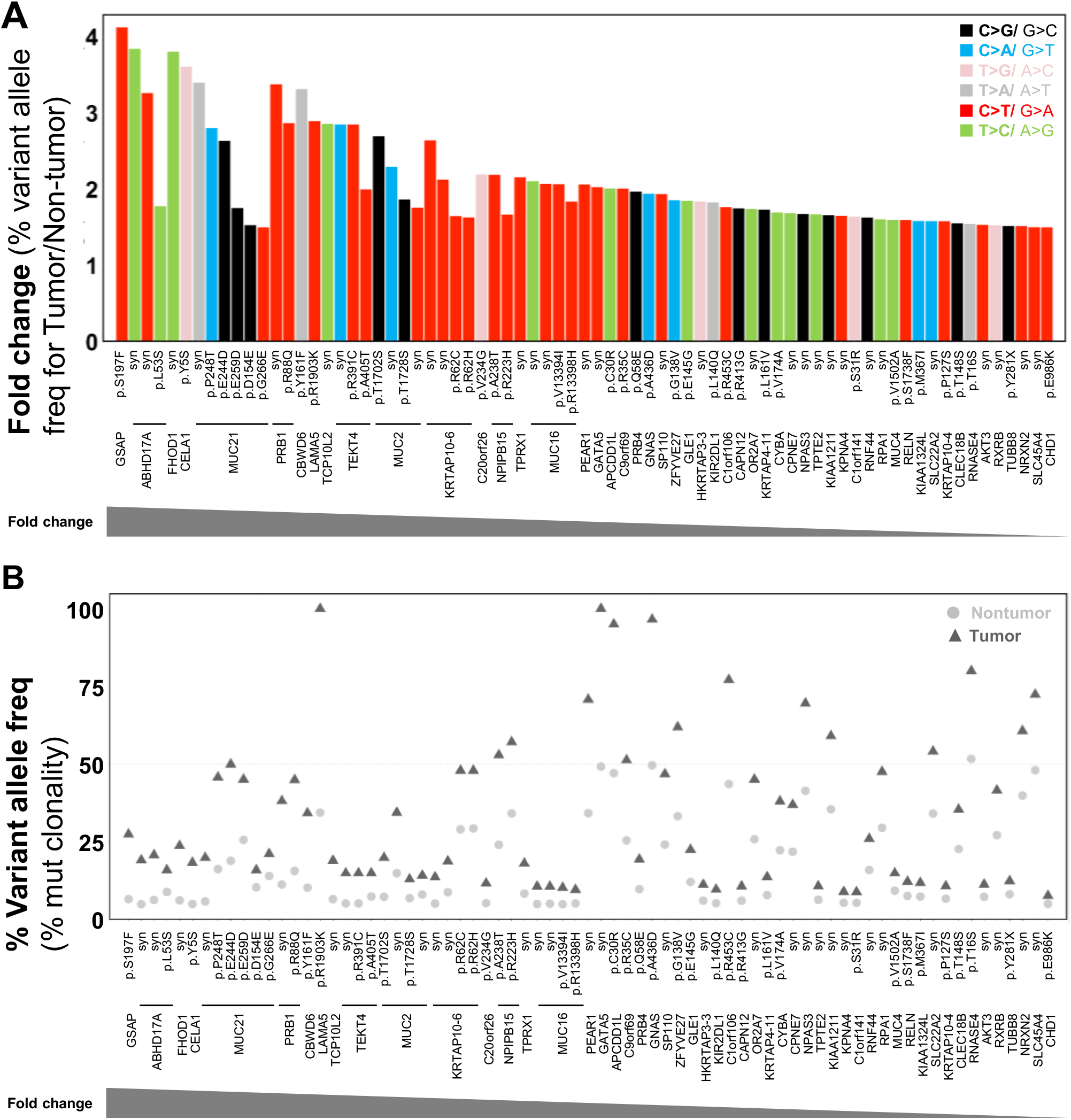
Somatic mutations that are clonally expanded during tumorigenesis were identified by whole exome sequencing. Mutations with variant allele frequencies (freq) less than 5% and mutations found in the matched normal cells (HME13) were removed. (A) Fold changes of clonally expanded somatic mutation allele frequencies between non-tumorigenic and tumorigenic cells colored by mutation types. (B) Variant allele frequencies (%) of each clonally expanded somatic mutation are displayed.

VAFs of 30 exonic mutations were expanded by two-fold or greater including G>T and C>G mutations in *MUC2*^[46,47]^, a C>T mutation in *LAMA5*^[48]^ (**Figure 3B, Table S1C**). Both genes are known to be associated with metastasis of breast cancer. Among the clonally expanded mutations by approximately two-fold (>1.9x), three heterozygous mutations in *APCDD1L, GATA5,* and *GNAS* with VAFs near 50% in non-tumorigenic cells became homozygous mutations with VAFs near 100% in tumorigenic cells (**Figure 3B, Table S1C**). *GATA5*^[49]^ and *GNAS*^[50,51]^ have been previously implicated in breast cancer. *APCDD1L* mRNA abundance has been associated with survival rates in other cancer types^[52]^ (**Table S1C**).

### 3.5. Significant differences in CNVs are observed between non-tumorigenic and tumorigenic cells

We determined somatic CNVs, as a percent of cell fraction, in non-tumorigenic *vs.* tumorigenic cells using CopyDetective^[34]^ with the WES data of the matched normal parental stem cells as the control (**Figure S2, Figure 4**). In order to identify true CNV regions that are significantly different between non-tumorigenic and tumorigenic cells, we filtered the raw CNV calls (**Figure S2**) to remove CNVs with less than ~5% cell fractions or CNVs with quality scores less than the default threshold (**Figure 4**). Statistically different changes (*p*<0.05) between the number of filtered raw CNVs in non-tumorigenic and tumorigenic cells were observed in 12 different chromosomes for duplications and in six chromosomes for deletions (**Figure 4**). Among these changes, the most statistically significant differences between non-tumorigenic and tumorigenic cells were found in chromosomes 5, 8, 16, and 22 for duplications and in chromosomes 8 and 15 for deletions (*p*<5×10^-4^) (**Figure 4C,D**). Specifically, non-tumorigenic cells have duplications in chromosomes 4p and 19p, while the tumorigenic cells have deletions in these locations (**Figure 4**). Tumorigenic cells have duplications or deletions in chromosomes 5, 8, and 14, and lost the deletion in chromosome 13q and duplication in chromosome 15 in comparison to the non-tumorigenic cells (**Figure 4**). The percent cell fractions of the filtered raw duplication calls were significantly higher in the tumorigenic cells in chromosomes 5, 8, and 16 (**Figure 4C**) than those in the non-tumorigenic cells. The percent cell fractions of the filtered raw deletion calls were significantly lower in chromosome 8 and higher in chromosome 15 of the tumorigenic cells (**Figure 4D**). Some CNVs are observed in both non-tumorigenic and tumorigenic cells, which include duplications in chromosomes 1q and 7q, and deletions in chromosome 3p, 8p, 15, and 22. Our results indicate that significant CNV differences are observed between non-tumorigenic cells and tumorigenic cells. These differences are more frequently found in duplications than in deletion.

**Figure 4.**
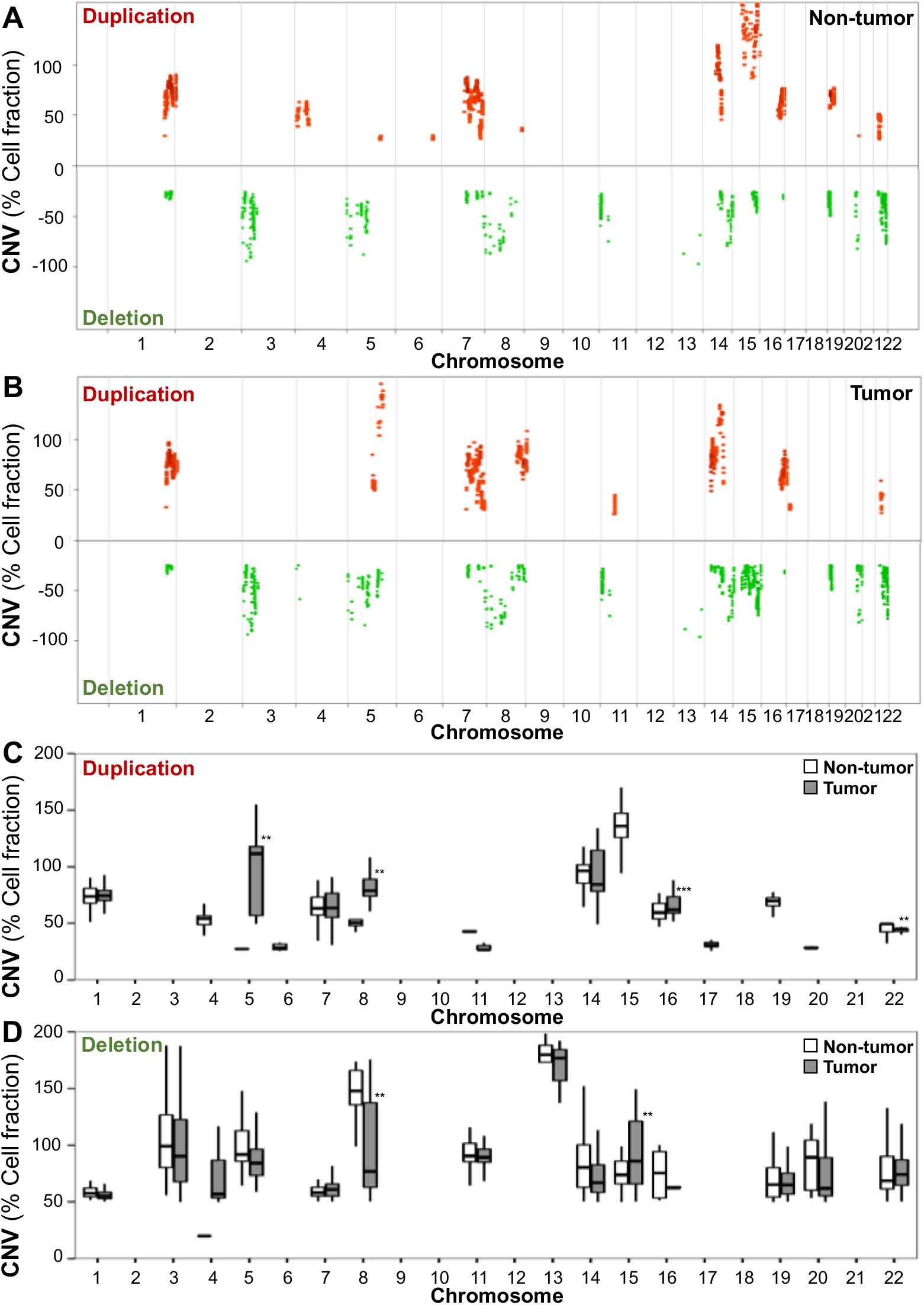
Significant differences in somatic CNVs are observed between human breast non-tumorigenic and tumorigenic cells. The fraction of cells with duplications and deletions that pass the default quality score filtration threshold of 10.76 in the **(A)** non-tumorigenic cells and **(B)** the tumorigenic cells were plotted for the 22 autosomal chromosomes. Duplications are represented in red with positive cell fractions and deletions are represented in green with negative cell fractions. The cell fractions of **(C)** duplications and **(D)** deletions in each chromosome are shown as box plots for non-tumorigenic and tumorigenic cells. The box plots represent the 75th and 25th percentile, with the middle line representing the median. The extreme line in the box plots represent 1.5 times interquartile range away from the upper and lower quartile. Significant differences in the cell fractions of duplications and deletions between non-tumorigenic and tumorigenic cells are indicated (**: *p*<5×10^-4^, ***: *p*<5×10^-10^) by the Mann-Whitney U test.

### 3.6. Cells align and migrate along ECM mimetic topography

We evaluated morphologies of non-tumorigenic and tumorigenic cells growing on nanopatterned (NP) and unpatterned (UP) surfaces. The results showed both cell types on NP surfaces aligned along the nano-grooves and displayed elongated morphologies, while these cells displayed no changes in cell morphologies on UP surfaces (**Figure 5**).

**Figure 5.**
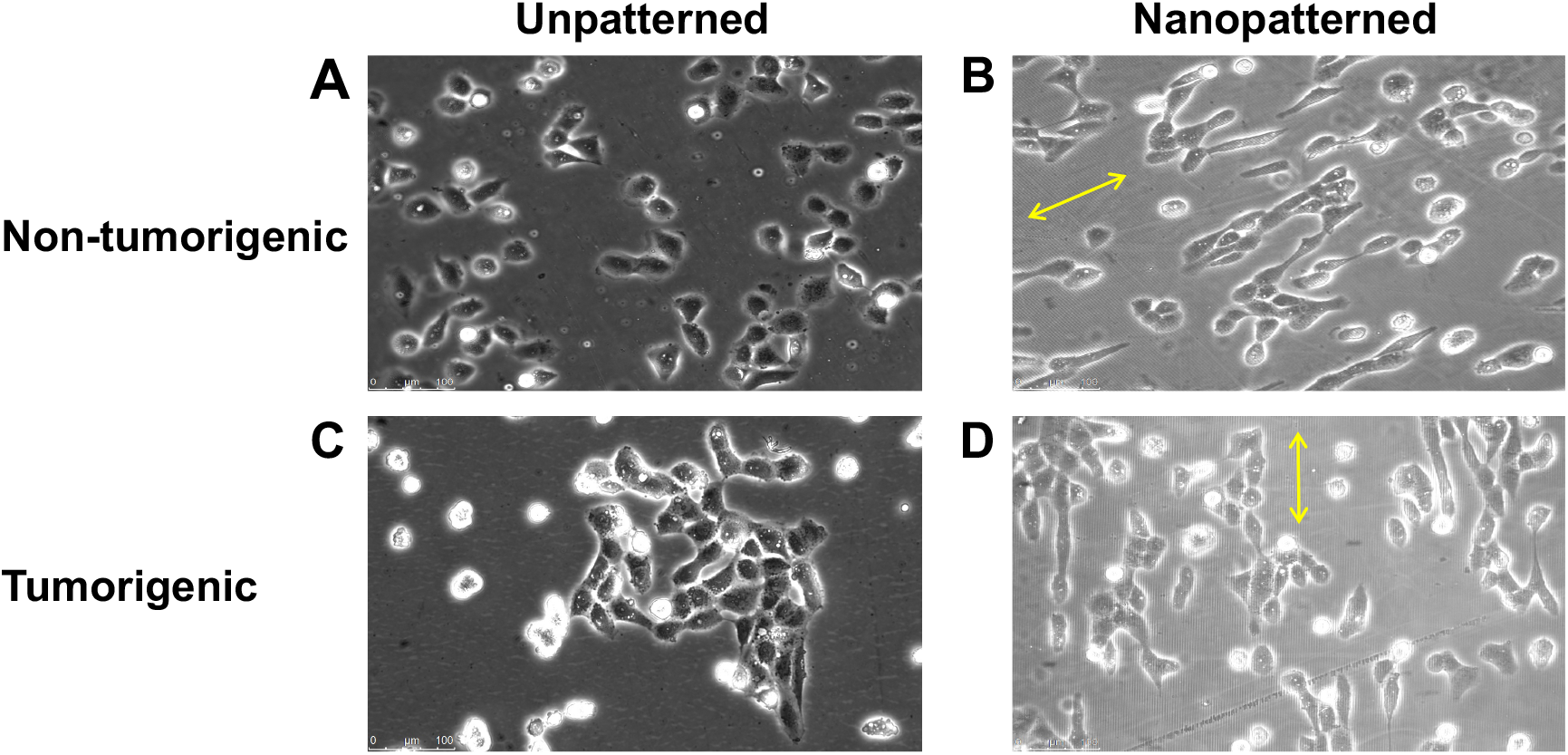
Both non-tumorigenic and tumorigenic human breast epithelial cells align along ECM mimetic topography. Human breast non-tumorigenic and tumorigenic cells were seeded on unpatterned and nanopatterned biocompatible polymer polyurethane-acrylate-301 (PUA301) substrates. Unlike the cells on unpatterned substrates **(A,C)**, both cell types align along the directions of nanogrooves and have elongated morphologies on nanopatterned substrates **(B,D)**. Images were taken 3.5 hours after cell seeding. Yellow arrows represent directions of nanogrooves. Scanning electron microscope image of a nanopatterned substrate is shown in **Fig S3**.

Single-cell migration trajectories were plotted in the *X* and *Y* directions with the cell’s start position set at coordinate (0,0). Trajectories of the non-tumorigenic or tumorigenic cells cultured on UP and NP surfaces were overlaid to produce a single-cell migration trajectory graph using an in-house MATLAB script. The trajectories of non-tumorigenic and tumorigenic cells on UP surfaces are observed to be nonlinear and multi-directional (**Figure 6A,C**). In contrast, the trajectories of both cell types on NP surfaces are mostly linear along the direction of the nano-groove (**Figure 6B,D**). Migration distances are also longer in cells on NP surfaces than cells on UP surfaces. Overall, the total distances migrated by the cells on NP surfaces are greater than those on UP surfaces for both cell types (**Figure 6**).

**Figure 6.**
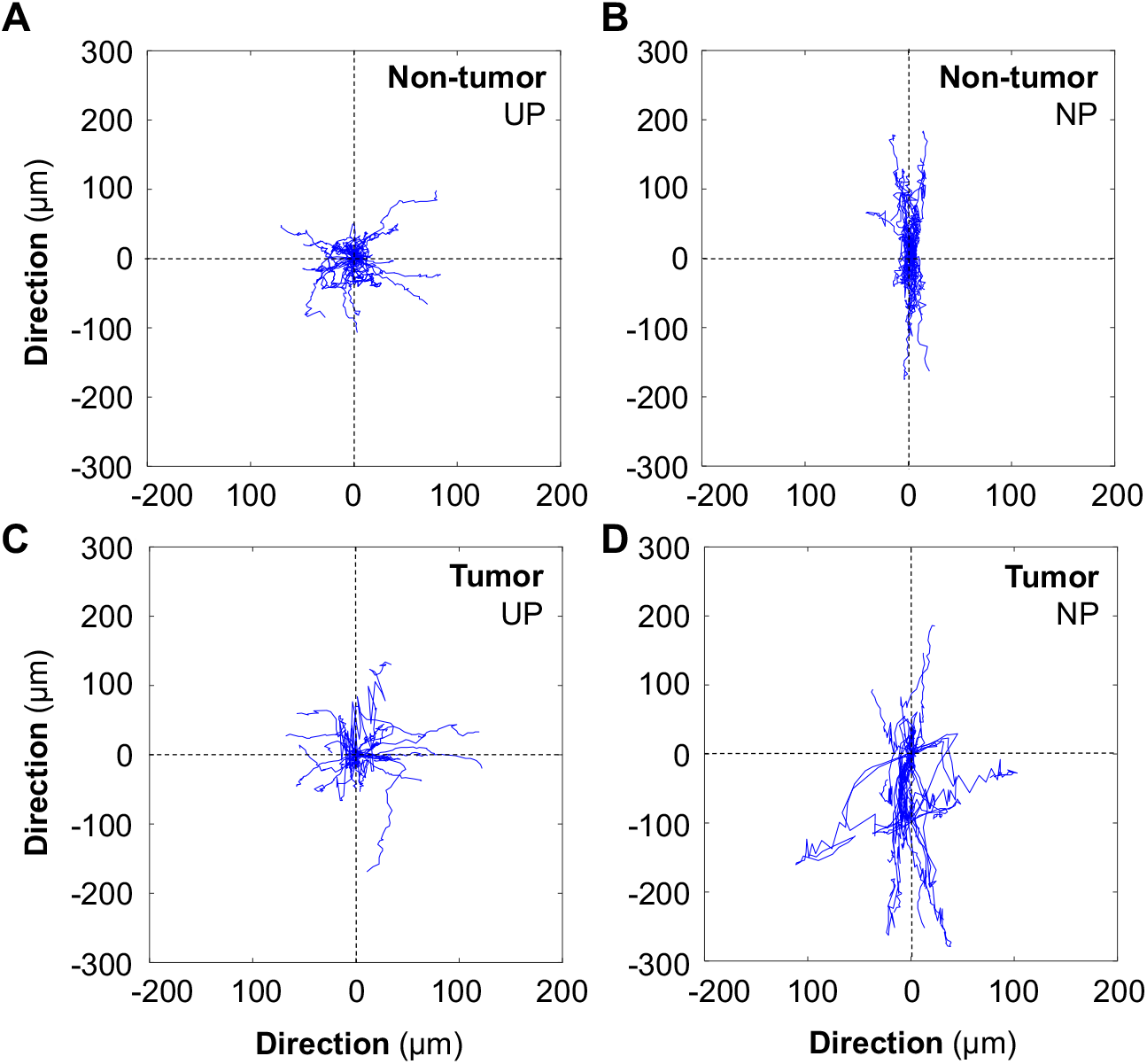
ECM-mimetic topography regulates the persistence of cell migration. The trajectories of a 12-hour single-cell migration are presented for human breast non-tumorigenic and tumorigenic cells cultured on unpatterned (UP) **(A,C)** and nanopatterned (NP) **(B,D)** PUA301 surfaces. Migration trajectories of each single-cell were overlaid for each group and all starting points were set to the coordinate (0,0).

### 3.7 ECM mimetic topography enhances the migration speed of tumorigenic cells but not non-tumorigenic cells

We analyzed the migration ability of non-tumorigenic and tumorigenic cells on different ECM topographical surfaces. The average single-cell migration speed of tumorigenic cells on NP surfaces was significantly higher (*p*<0.05) than those of tumorigenic cells on UP surfaces and non-tumorigenic cells on NP and UP surfaces (**Figure 7**). In contrast, no significant difference was observed between the average migration speeds of tumorigenic (9.88 μm/hr) and non-tumorigenic (9.44 μm/hr) cells on UP surfaces (**Figure 7**). This indicates that ECM-mimetic topography enhances migration speed of tumorigenic cells but not non-tumorigenic cells. The enhanced migratory ability of tumorigenic cells is clearly lost in conventional two-dimension culture conditions, demonstrating they lack the capacity to recapitulate behaviors of tumors cell observed *in vivo.*

**Figure 7.**
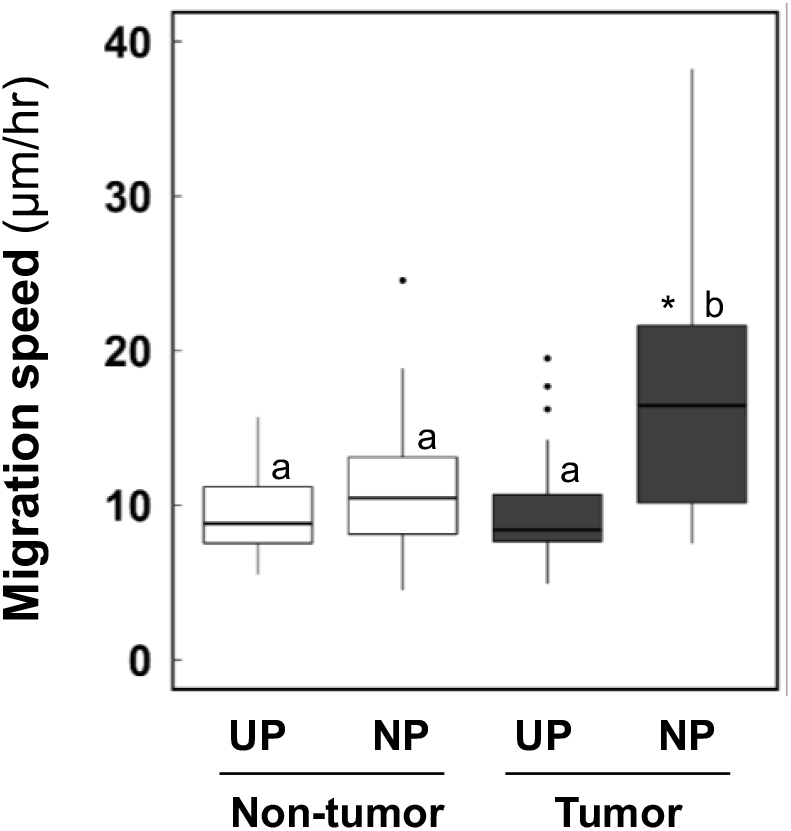
ECM-mimetic topography enhances the migration speed of tumorigenic cells but not non-tumorigenic cells. Non-tumorigenic and tumorigenic cells were seeded on unpatterned (UP) and nanopatterned (NP) PUA301 substrates. Single-cells were individually tracked to calculate migration speed (non-tumorigenic UP n=40, non-tumorigenic NP n=50, tumorigenic UP n=26, tumorigenic NP n=28). Box and whisker plots show an increase in the migration speed of tumorigenic cells on NP (ECM mimetic topography) (a,b: *p*<0.05 by the ANOVA on ranks). Significant differences in distribution of individual single-cell migration speed are indicated (*: *p*<0.05) by the Kolmogorov-Smirnov test.

A greater range of individual single-cell migration speeds was observed in tumorigenic cells (NP 8-38 μm/hr; UP 5-20 μm/hr) than in non-tumorigenic cells (NP 5-25 μm/hr; UP 6-16 μm/hr) (**Figure 7**). This wider range of single-cell migration speeds reflects heterogeneity of tumorigenic cells which contain a significantly higher subpopulation of CSCs than non-tumorigenic cells (**Figure 1**).

An independent cell culture experiment was conducted for the single-cell migration analysis with tumorigenic cells. All experimental conditions were maintained, and cell migration was quantified in the same manner as for the results presented in Figures 5–7. Consistent results were obtained. For example, the average migration speed was significantly higher for tumorigenic cells on NP surface than on UP surface (*p* < 0.001) (**Figure S4**). Furthermore, a similar significant difference in the distribution of individual cell migration speeds and migration trajectories (**Figure S5**) of tumorigenic cells was observed between UP and NP surfaces.

## 4. Discussion

In this study, we evaluated the cellular migration of human breast CSCs by quantifying and comparing migratory properties of non-tumorigenic and tumorigenic cells. Both cell types were sequentially derived from the same parental normal human breast epithelial stem cells (HBECs) with stem cell features^[18,22,23]^. To recapitulate tumor microenvironment, we cultured these cells (non-tumorigenic vs tumorigenic) on our ECM-mimetic platforms. Our results demonstrate that tumorigenic cells lose their enhanced migratory and heterogeneous properties of cellular sensing on surfaces lacking ECM-mimetic structures. This loss of increased cellular migration and heterogeneity of tumorigenic cells is recovered on an ECM-mimetic surface.

WES results reveal an increased somatic mutation burden in tumorigenic cells than in non-tumorigenic cells. A greater mutational burden has been associated with metastatic breast cancer^[53]^. With respect to the distribution of synonymous and nonsynonymous somatic mutations, the proportion of synonymous mutations was greater than that of nonsynonymous mutations in tumorigenic cells. The first evidence that not all synonymous mutations are silent mutations was manifested by Kimchi et. al., which found that a synonymous mutation in the multidrug resistance 1 (*MDR1*) gene altered the effectiveness of inhibitors of P-glycoprotein^[54]^. Since then, synonymous mutations have been receiving more attention insofar as their potential impacts in cancer development. For example, some synonymous mutations in oncogenes can act as driver mutations by altering exonic motifs^[55]^. Synonymous mutations can alter splicing sites, mRNA stability, miRNA binding sites or translation and can lead to exon skipping and thus a change in protein structure of some of tumor suppressor genes/proteins^[56]^. In a pan-cancer analysis of 659,194 synonymous mutations identified in TCGA, ICGC, and COSMIC databases, 176,594 synonymous mutations were found recurrently across all tumors.

We identified genes that were mutated exclusively in tumorigenic cells or clonally expanded during breast tumorigenesis. These genes can be candidate genes for breast tumorigenesis and metastasis. Among the 81 genes that contain mutations exclusive to tumorigenic cells, *ERBB2*^[57]^, *FANCD2*^[38,39]^, *MMP13*^[40]^, *MSH2*^[41]^, *RET*^[42]^, *ITCH*^[43]^, *CHD8*^[44]^, and *SREBP-2*^[45]^ have been previously reported to be associated with increased risk of breast cancer. It was demonstrated that the overall survival of patients of breast tumors containing *ERBB2* mutations was lower than that of patients with wild type *ERBB2*^[57]^. Other genes in the list include *CLASP1, RAP2B, RET,* and *SEMA3E,* all of which have a role in regulating cell migration or cell-matrix adhesion based on the Gene Ontology^[58]^. *RAP2B*, in particular, has been previously reported to be overexpressed in multiple breast cancer cell lines and to increase calcium levels^[59]^. The elevated calcium level promotes breast cancer cell migration and invasion via the calcium related ERK1/2 signaling pathway^[59]^. Some of clonally expanded mutations in tumorigenic cells were found in genes that have been associated with breast cancer, such as *AKT3*^[60]^, *GNAS*^[50,51]^, *GATA5*^[49]^, *LAMA5*^[48]^, *MUC2*^[46,47]^ *NRXN2*^[36]^, *RELN*^[61]^, and *RPA1*^[62]^. Notably, mutations in *LAMA5* and *MUC2* have high fold-changes of 2.9 and 2.7, respectively. A previous study reported that exosomes derived from non-metastatic breast cancer cells compared to those derived from metastatic breast cancer cells were differentially enriched for *LAMA5,* a gene implicated for cell adhesion^[48]^. *MUC2* mediated proliferation and metastasis of breast cancer cells^[46,47]^. Expression of *MUC2* was weak or absent in healthy breast tissue, and patients with high *MUC2* expression were observed to have a significantly shorter survival^[46]^. In addition, mutations in *GNAS* and *GATA5* expanded from heterozygous mutations (VAFs near 50%) to homozygous mutations (VAFs near 100%), with the mutation in *GNAS* having the highest CHASMplus scores among clonally expanded mutations by greater than 1.5-fold. A previous study reported that the high expression of *GNAS* is associated with distal metastasis and poor survival rates based on 150 breast cancer samples^[51]^. siRNA-mediated *GNAS* knockdown experiments showed reduced viability and proliferation, impaired migration ability, and induced G1 cell cycle arrest in breast cancer cells^[50,51]^. Potential mechanisms for these changes induced by the high expression of *GNAS* involve the activation of the PI3K/AKT signaling pathway^[51]^ and overexpression of the extra-long Gas (XLas) isoform, which induces cAMP levels^[50]^. *GATA5* has been reported to activate the progesterone receptor gene promoter under the presence of the progesterone receptor +331G/A gene variant, a variant that is associated with breast cancer risk^[49]^. Furthermore, *GATA5* has been previously investigated as a potential tumor suppressor in colorectal cancer^[63]^, hepatocellular carcinoma^[64]^, and lung adenocarcinoma^[65]^. These mutations, acquired or expanded during breast tumorigenesis, that we identified, can account for, at least in part, enhanced migratory ability of tumorigenic cells.

CNVs in specific regions of chromosomes can be involved in tumor invasion and migration^[14,66]^. We examined somatic CNV profiles of non-tumorigenic and tumorigenic cells and identified genomic regions that are significantly different in CNVs between non-tumorigenic and tumorigenic cells. These identified regions were reported to be associated with breast cancer cell invasiveness^[67,68]^. For example, a loss of heterozygosity (LOH) of the chromosome 8p region increased motility and invasiveness in MCF10A breast cancer cells^[67]^. A previous study by *Pariyar et al.* reported that amplifications of the chromosome regions 1q, 8q, 19p and 19q, 2p, 5p and the deletions in the chromosome regions 8p, 5q, and 19p were common in invasive ductal carcinomas (IDCs) of triple negative breast cancer tissue specimens^[68]^. Our results showed that the tumorigenic cells gained significantly higher number of duplications in the chromosome region 8q and 5p but lost the duplications in 19p. Both tumorigenic and non-tumorigenic cells had deletions in the chromosome region 8p, 5q, and 19p. Overall, our results indicate that the tumorigenic cells have distinct somatic CNVs, which could contribute to their enhanced invasiveness.

Collagen I fibers have been used in tumor models to simulate the interaction between cancer cells and the tumor microenvironment, such as invasion^[7]^ and intravasation^[69]^. To recapitulate tumor microenvironment, we have applied an ECM-mimetic, nanotopographically-defined cell culture platform. In addition, we have integrated the advantages of highly controllable topography on quasi-3D nanopatterned substrates as the mechanical nanoscale contact guidance cue and collagen I as the biochemical cue. Previously, we applied this platform to study cell migration in human brain cancer^[10]^, melanoma^[70]^, and breast immortalized^[11]^ cells. Features of our ECM-mimetic culture platform include treatment of the surface with collagen type I to enhance cellular attachment^[71]^ and 800 nm spacing of nanogrooves to resemble fibril spacings in mammary collagen^[11,72]^. We have demonstrated that this platform provides nanoscale contact guidance cues evidenced by cell alignment along edges of nanogrooves and enhancement in cell migration. Contact guidance is a strong regulator of directed migration^[8]^ and plays a major role in cancer cell invasion^[73]^. The presence of nanoscale contact guidance in our study is manifested by the aligned and elongated morphologies of both non-tumorigenic and tumorigenic cells along the edges of nanogrooves.

We observed that the average migration speed of tumorigenic cells is significantly higher on nanopatterned surfaces compared to those on unpatterned surfaces, suggesting that nanoscale contact guidance cues enhance the cellular migration of breast tumorigenic cells. In contrast, we observed no change in migration speed for non-tumorigenic breast cells on nanopatterned surfaces *vs.* unpatterned surfaces. This indicates that contact guidance cues alone cannot enhance the migratory ability of non-tumorigenic cells. It is possible that non-tumorigenic cells may lack mutations and epigenetic changes that are present in tumorigenic cells that induce more responsiveness to contact guidance cues. Selective enhancement of migratory ability for tumorigenic cells on our ECM-memetic topography supports the idea that this platform can recapitulate cellular migration observed in a tumor microenvironment fostered by the ECM structure.

The linear, as opposed to random, nature of cell trajectories in the presence of nanoscale guidance cues has been suggested as a possible mechanism for enhanced cell migration speed^[11,74,75]^. Our results indicate that trajectories for both non-tumorigenic and tumorigenic cells are more linear on nanopatterned surfaces than those on the unpatterned surfaces; however, increased average migration speed was only observed for tumorigenic cells. This suggests that contact guidance cues likely trigger distinct alterations in cells that enhance their physical movement rather than through more directed and linear migration.

Other potential mechanisms of the enhanced migration of tumorigenic cells on the ECM-mimetic platform can be related to cell types. For example, *Ray et al.* demonstrated that aligned topography promoted migration speed through contact guidance regulated by focal adhesion maturation and F-actin alignment^[8]^. Mesenchymal-type cancer cells display a higher sensitivity to the contact guidance of the aligned topography compared to epithelial type cancer cells. A study on the migration of cancer cells in 3D collagen fibers reported that aligned collagen matrices enhanced motility in cancer stem cells (CSCs)^[75]^. Biophysical analyses of cell migration indicate that smaller cell sizes and higher phenotypic plasticity and protrusive activity may be the cause of the increased migration speed of CSCs. *Domura et al.* reported that the epithelial-mesenchymal transition (EMT) of cancer cells has been reported on aligned topography, suggesting that EMT may influence cancer cell migration^[76]^.

Cell migration can be also influenced by biomolecular and biophysical properties of the cell. Expression levels of cell surface receptors, such as receptor tyrosine kinases, transmit signals from the extracellular environment to modulate cell migratory behavior. Within the same cell type, receptor expression levels can be heterogenous in cancer tumors. *Smith et al.* showed that subpopulations of glioblastoma cells with high PDGF receptor alpha (PDGFRα) had a correlation with enhanced migration on nanopatterned surfaces; however, the authors noted that fidelity of the receptor and downstream effectors in this pathway likely account for differences among individual cells in their assay^[10]^. Actomyosin complexes downstream of receptor tyrosine kinases promote protrusions that propel migratory cells. In 3D microenvironments, protrusive pseudopodial structures have been observed even in the inhibition of actomyosin contractility, suggesting the role of biophysical properties of the cell membrane and proteins associated with focal adhesions to promote enhanced migration of cancer cells ^[77]^.

In addition to higher average migration speeds on NP *vs.* UP surfaces, tumorigenic cells displayed significant differences in the distributions of individual cell speeds on NP surfaces than on UP surfaces, as verified by the Kolmogorov-Smirnov test. This suggests heterogeneity in tumorigenic cells. No such significant differences in individual cell migration speeds were observed for both cell types on unpatterned surfaces and for non-tumorigenic cells on nanopatterned surfaces. To our knowledge, this migration aspect has not yet been investigated at single-cell level, as other studies focused on collective cell migration. Individual breast tumorigenic cells possibly acquire different somatic mutations and epigenetic alterations that change how each cell senses and responds to contact guidance cues in the tumor microenvironment. Future studies can characterize various cell migration speed subpopulations of tumorigenic cells on nanopattern surfaces to ascertain, for example, if higher speeds are found in breast CSC-like populations. This would test if CSCs are more susceptible to the contact guidance cues associated with enhanced migration.

In the present study, we have investigated mutational changes and copy number variations (CNVs) of the whole exome and performed a single-cell migration assessment on the ECM-mimetic nanoscale cell culture platform in tumorigenic *vs*. non-tumorigenic cells (both derived from the same normal parental stem cells). We have identified regions of CNVs and groups of mutations as candidates associated with breast tumorigenesis and the varying degrees of breast tumor invasiveness. Our data indicate that ECM-mimetic topography selectively enhances migratory ability of breast tumorigenic cells, but not non-tumorigenic cells. This demonstrates that our nanotopographically defined platform can recapitulate an ECM structure of the tumor microenvironment to study cell migration.

## Author Contributions

Conceptualization: EHA; Experiments: EHA, JYK, SHTL, PK; Data curation: EHA, SHTL, JYK, PK, ZD; Data analysis: SHTL, EHA, JYK, ZD, PK; Funding acquisition: EHA; Investigation: EHA, SHTL, JYK, PK, ZD; Methodology: SHTL, EHA, JYK, ZD, PK; Project administration, EHA; Resources: EHA; Supervision: EHA; Visualization: SHTL, EHA, JYK, ZD, CYS, PK; Writing: SHTL, EHA, JYK; Editing and review: SHTL, EHA, ZD, CYS.

## Funding

The research was supported by grants from the National Institute of Environmental Health Sciences (NIEHS) P30 ES007033 sponsored-University of Washington (UW) Center for Exposures, Diseases, Genomics and Environment (EDGE) pilot grant (to EHA), UW Office of Research Royalty Research Fund (to EHA), and the National Cancer Institute (NCI) P30 CA015704-39 Fred Hutchinson Cancer Research Center-UW Cancer Consortium Support Grant (to EHA), NCI R21 CA220111 (to EHA) of the National Institutes of Health (NIH). The content is solely the responsibility of the authors and does not necessarily represent the official views of the NIH.

## Acknowledgments

We thank Drs. Chia-Cheng Chang, Brad L. Upham, and James E. Trosko for obtaining non-tumorigenic immortalized and tumorigenic/xenograft cells; Dr. Deok-Ho Kim for providing access to Nikon Eclipse Ti microscope and Femto Science SUTE-MPR Plasma system; Howard Nebeck for editing and proofreading the manuscript; Jiwon Min for managing references of the manuscript, and Dr. Allister G. Suarez for comments on the manuscript.

## Notes

### Competing Interest Statement

The authors have declared no competing interest.

### Summary of Updates

Copy number variation (CNV) analysis results have been added to the manuscript.

