## Supplementary Information for "Changes of Mutations and Copy-number and Enhanced Cell Migration during Breast Tumorigenesis"

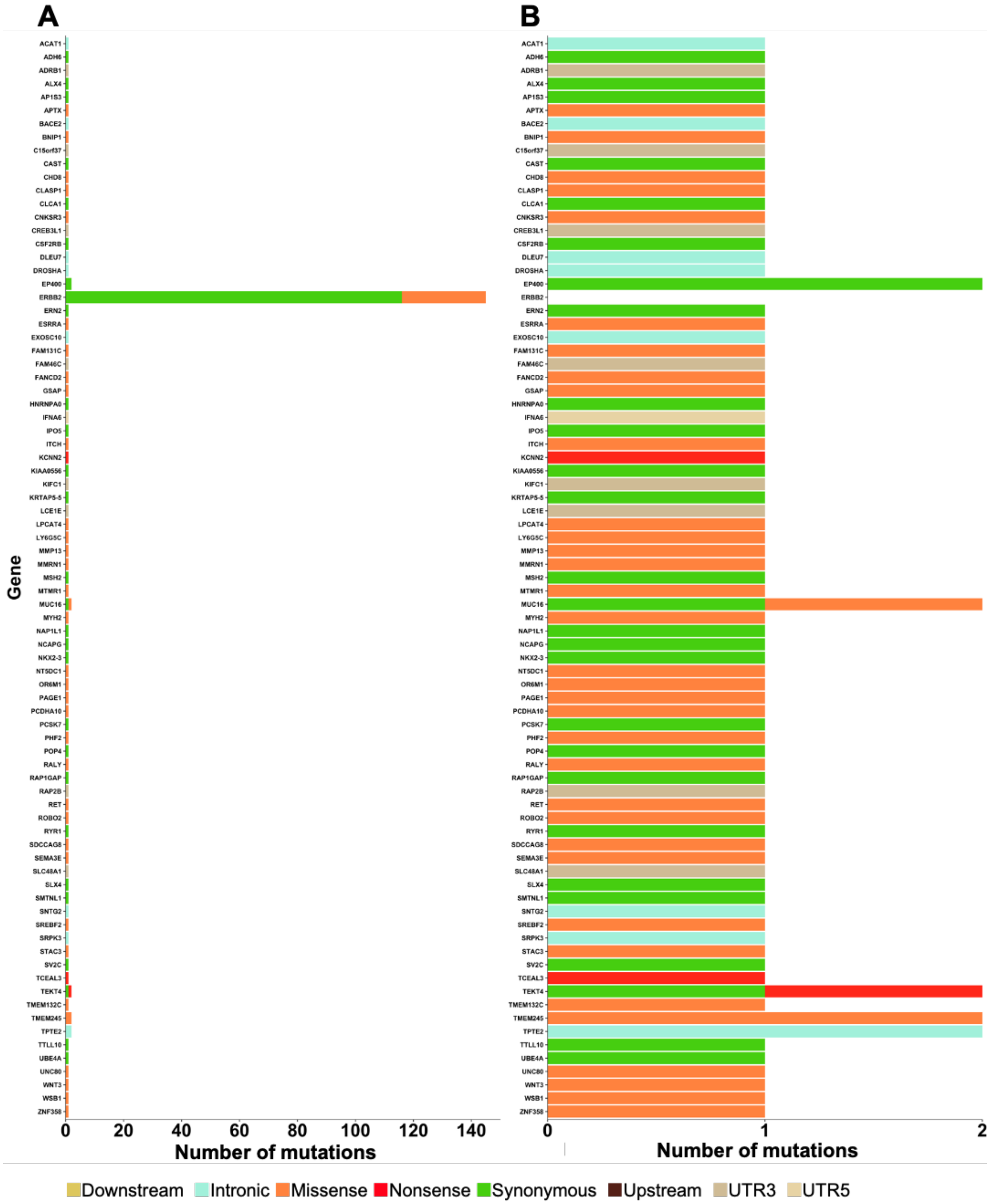


**Figure S1.** Numbers of somatic mutations exclusively found in tumorigenic cells for each gene including **(A)** and excluding **(B)** *ERBB2* gene. Mutations with variant allele frequencies less than 5% and mutations found in the matched normal cells (HME13) were removed. Colors represent proportion of each mutation type. Abbreviations used are: UTR3, three prime untranslated region; UTR5, five prime untranslated region.

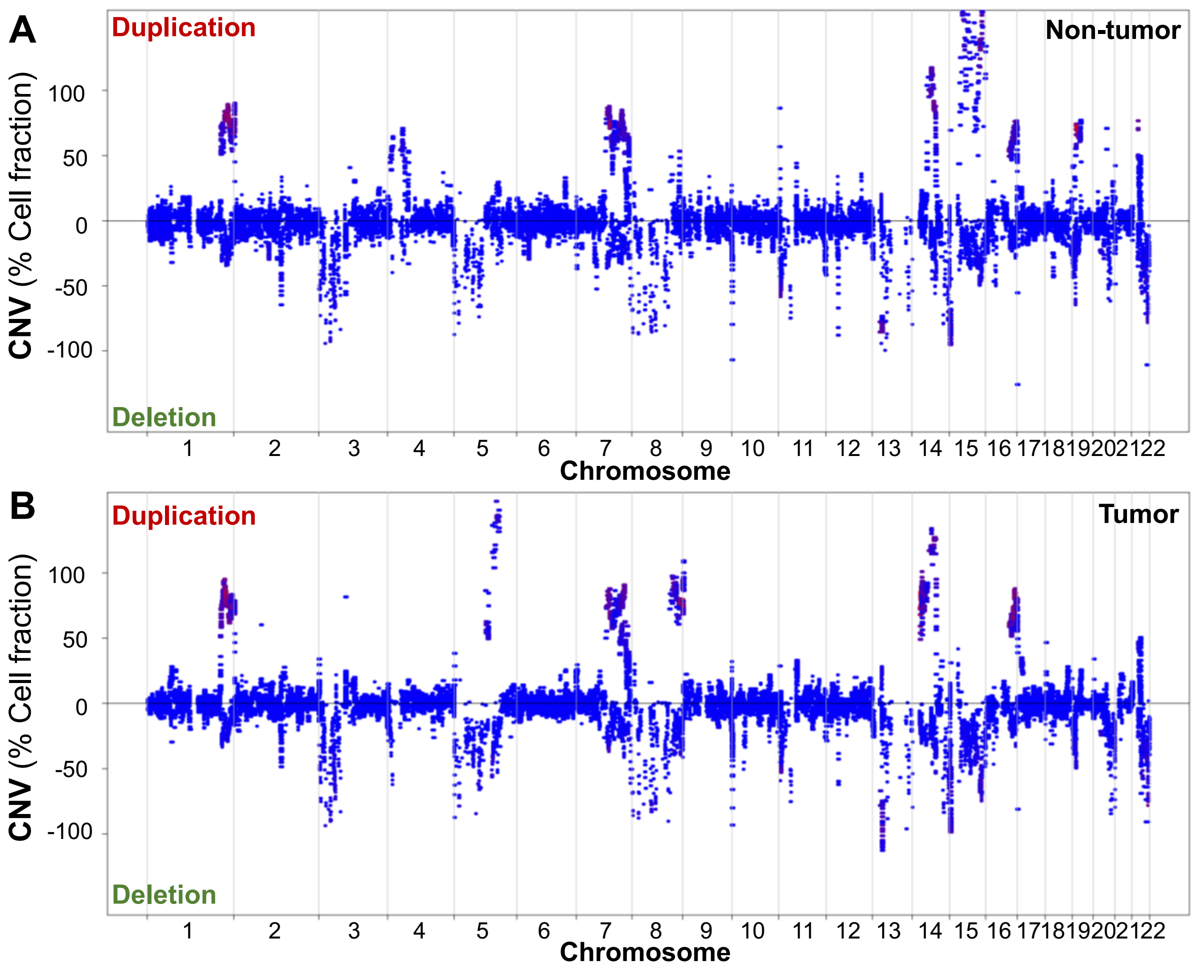


**Figure S2.** The fraction of raw copy number variants (CNVs) in the **(A)** non-tumorigenic cells and **(B)** the tumorigenic cells were plotted for the 22 autosomal chromosomes. Duplications are represented with positive cell fractions and deletions are represented with negative cell fractions.

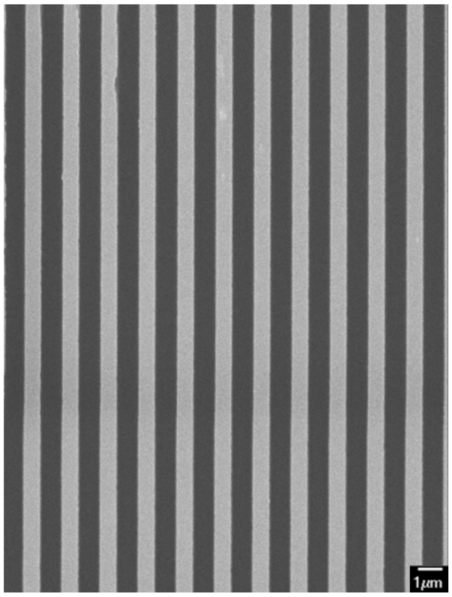


**Figure S3**. A representative scanning electron microscope image of extracellular matrix (ECM)-mimetic nanotopographical model. Each nano-groove is 800nm by 800nm and are separated by 800nm.

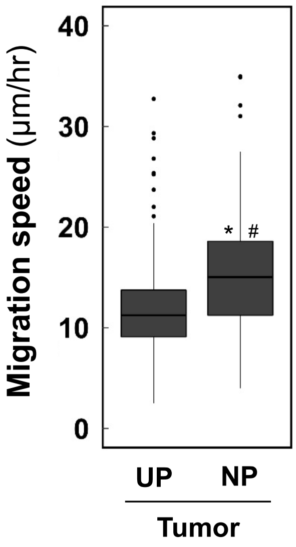


**Figure S4.** Independent cell culture experiments generate reproducible results of migration speeds of human breast tumorigenic cells. Single-cells were individually tracked to calculate the speed of tumorigenic cell migration on unpatterned (UP) and nanopatterned (NP) biocompatible polymer polyurethane-acrylate-301 substrates (tumorigenic UP n=77 and tumorigenic NP n=76). Box plots indicate average migration speed of each group and distributions of individual cell migration speeds. Significant differences in average migration speeds between the two groups are indicated ($^{*}$: $p<0.001$) by the Mann-Whitney Rank Sum test. Significant difference in distribution of individual cell migration speeds is indicated ($^{\#}$: $p<0.05$) by the Kolmogorov-Smirnov test.

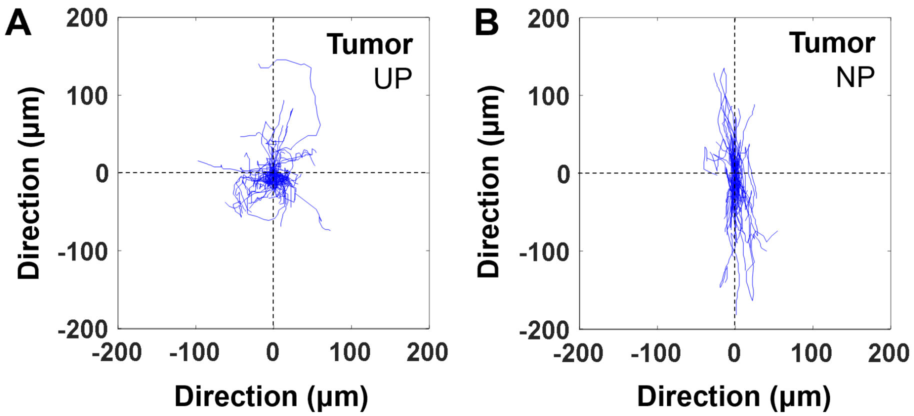


**Figure S5.** Independent cell culture experiments generate reproducible results for trajectories of a 12-hour single-cell migration in human breast tumorigenic cells cultured on unpatterned (UP) **(A)** and nanopatterned (NP) **(B)** biocompatible polymer polyurethane-acrylate-301 substrates. Migration trajectories of each cell were overlaid for each group and all starting points were set to the coordinate (0,0).

**Table S1**

**[A]** Somatic mutations exclusive to immortalized cells were identified by whole exome sequencing. Mutations with variant allele frequencies less than 5% were removed. CHASMplus scores were calculated based on all 32 cancer types and on breast cancer alone.

| Chr | Position | Reference Allele | Alternate Allele | Mutation Consequence | Gene symbol | Amino Acid change | CHASMplus | CHASMplus_BRCA |
|---|---|---|---|---|---|---|---|---|
| 1 | 11839124 | A | C | missense_variant | C1orf167 | T904P | 0.018 | 0.03 |
| 1 | 12853509 | A | C | missense_variant | PRAMEF1 | S45R | 0.002 | 0.001 |
| 1 | 12980232 | G | A | stop_retained_variant | PRAMEF7 | *475 | | |
| 1 | 16262471 | A | C | missense_variant | SPEN | T3246P | 0.171 | 0.228 |
| 1 | 21016827 | T | C | missense_variant,splice_region_variant | KIF17 | E412G | 0.027 | 0.02 |
| 1 | 39879403 | T | C | missense_variant | KIAA0754 | S1020P | 0.055 | 0.042 |
| 1 | 65830355 | A | G | synonymous_variant | DNAJC6 | syn | | |
| 1 | 152186042 | A | G | missense_variant | HRNR | L2688S | 0.026 | 0.02 |
| 1 | 152770446 | A | G | missense_variant | LCE1D | N59S | 0.006 | 0.017 |
| 1 | 152883134 | A | G | synonymous_variant | IVL | syn | | |
| 1 | 152883164 | T | G | missense_variant | IVL | D297E | 0.011 | 0.007 |
| 1 | 167385267 | T | A | 3_prime_UTR_variant | POU2F1 | -- | | |
| 1 | 175116119 | A | G | missense_variant | TNN | E1271G | 0.02 | 0.009 |
| 1 | 186276121 | A | C | missense_variant | PRG4 | T424P | 0.01 | 0.007 |
| 1 | 186276291 | A | G | synonymous_variant | PRG4 | syn | | |
| 1 | 240370967 | C | G | missense_variant | FMN2 | A952G | 0.014 | 0.017 |
| 1 | 240371463 | C | G | synonymous_variant | FMN2 | syn | | |
| 1 | 244219959 | T | C | 3_prime_UTR_variant | ZBTB18 | -- | | |
| 2 | 37276942 | A | T | synonymous_variant | HEATR5B | syn | | |
| 2 | 37276957 | T | G | synonymous_variant | HEATR5B | syn | | |
| 2 | 37276964 | G | A | missense_variant | HEATR5B | T843I | 0.096 | 0.057 |
| 2 | 101521208 | G | A | 5_prime_UTR_variant | NPAS2 | -- | | |
| 2 | 234178652 | C | G | missense_variant | ATG16L1 | R216G | 0.137 | 0.055 |
| 3 | 58109406 | T | A | missense_variant | FLNB | I1238K | 0.25 | 0.157 |
| 3 | 58109408 | G | A | missense_variant | FLNB | E1239K | 0.261 | 0.164 |
| 3 | 75786315 | C | T | missense_variant | ZNF717 | R820K | 0.007 | 0.023 |
| 3 | 183888392 | T | C | missense_variant | DVL3 | L667P | 0.192 | 0.093 |
| 3 | 195505766 | T | C | missense_variant | MUC4 | S4229G | 0.032 | 0.022 |
| 3 | 195506022 | C | G | synonymous_variant | MUC4 | syn | | |
| 3 | 195506254 | T | G | missense_variant | MUC4 | H4066P | 0.037 | 0.027 |
| 3 | 195506332 | G | A | missense_variant | MUC4 | S4040L | 0.045 | 0.029 |
| 3 | 195506692 | C | G | missense_variant | MUC4 | R3920P | 0.043 | 0.027 |
| 3 | 195513934 | T | G | missense_variant | MUC4 | H1506P | 0.039 | 0.028 |
| 3 | 195513942 | A | G | synonymous_variant | MUC4 | syn | | |
| 3 | 195514903 | G | A | missense_variant | MUC4 | T1183M | 0.032 | 0.021 |
| 4 | 106163131 | C | A | intron_variant | TET2 | -- | | |
| 4 | 123128697 | G | T | splice_region_variant,synonymous_variant | KIAA1109 | syn | | |
| 4 | 152583882 | T | G | synonymous_variant | FAM160A1 | syn | | |
| 4 | 152583895 | T | C | missense_variant | FAM160A1 | S1038P | 0.006 | 0.006 |
| 4 | 153864389 | A | C | synonymous_variant | FHDC1 | syn | | |
| 4 | 153896049 | A | C | missense_variant | FHDC1 | T536P | 0.008 | 0.013 |
| 4 | 183522111 | G | C | synonymous_variant | TENM3 | syn | | |
| 5 | 843745 | C | T | missense_variant | ZDHHC11 | G200R | 0.016 | 0.017 |
| 5 | 78610466 | T | C | synonymous_variant | JMY | syn | | |
| 5 | 87516527 | T | A | missense_variant | TMEM161B | Y100F | 0.05 | 0.016 |
| 6 | 30672200 | A | G | missense_variant | MDC1 | L1587P | 0.084 | 0.073 |
| 6 | 30954298 | G | A | missense_variant | MUC21 | G116R | 0.029 | 0.019 |
| 6 | 30954378 | A | G | synonymous_variant | MUC21 | syn | | |
| 6 | 30955160 | G | A | missense_variant | MUC21 | S403N | 0.028 | 0.018 |
| 6 | 31322910 | G | A | missense_variant | HLA-B | A329V | 0.223 | 0.03 |
| 6 | 31322915 | G | C | synonymous_variant | HLA-B | syn | | |
| 6 | 32609212 | C | T | missense_variant | HLA-DQA1 | R70W | 0.065 | 0.049 |
| 6 | 41554882 | A | C | missense_variant | FOXP4 | T216P | 0.129 | 0.167 |
| 6 | 50681806 | A | G | missense_variant,splice_region_variant | TFAP2D | E13G | 0.008 | 0.004 |
| 6 | 109906342 | C | T | missense_variant | AK9 | E700K | 0.011 | 0.026 |
| 7 | 29923929 | G | C | synonymous_variant | WIPF3 | syn | | |
| 7 | 73011713 | T | G | missense_variant | MLXIPL | T468P | 0.022 | 0.022 |
| 7 | 82784441 | A | G | missense_variant | PCLO | S506P | 0.031 | 0.044 |
| 7 | 82784463 | C | T | synonymous_variant | PCLO | syn | | |
| 7 | 82784493 | C | T | synonymous_variant | PCLO | syn | | |
| 7 | 99764683 | G | T | synonymous_variant | GAL3ST4 | syn | | |
| 7 | 103202356 | A | C | missense_variant | RELN | V1752G | 0.033 | 0.022 |
| 7 | 123302942 | T | C | synonymous_variant | LMOD2 | syn | | |
| 8 | 623613 | G | C | missense_variant | ERICH1 | Q247E | 0.004 | 0.005 |
| 8 | 10465245 | A | T | synonymous_variant | RP1L1 | syn | | |
| 8 | 87680396 | G | T | missense_variant,splice_region_variant | CNGB3 | A165E | 0.007 | 0.005 |
| 8 | 90796352 | T | A | missense_variant | RIPK2 | N338K | 0.016 | 0.009 |
| 9 | 21968204 | T | G | 3_prime_UTR_variant | CDKN2A | -- | | |
| 9 | 38398289 | G | A | 3_prime_UTR_variant | ALDH1B1 | -- | | |

| | | | | | | | | |
|---|---|---|---|---|---|---|---|---|
| 9 | 46386764 | T | C | missense_variant | FAM27E1 | D80G | | |
| 9 | 70919114 | C | G | missense_variant | FOXD4L3 | P416R | 0.004 | 0.002 |
| 9 | 73479364 | T | A | missense_variant | TRPM3 | T81S | 0.03 | 0.024 |
| 9 | 102590828 | A | C | synonymous_variant | NR4A3 | syn | | |
| 9 | 112185006 | A | G | synonymous_variant | PTPN3 | syn | | |
| 9 | 139342874 | C | A | intron_variant | SEC16A | -- | 0.078 | 0.05 |
| 10 | 121356005 | C | A | 5_prime_UTR_variant | TIAL1 | -- | | |
| 11 | 1629277 | A | G | synonymous_variant | KRTAP5-3 | syn | | |
| 11 | 1651393 | C | G | missense_variant | KRTAP5-5 | A108G | 0.009 | 0.017 |
| 11 | 49175793 | G | A | synonymous_variant | FOLH1 | syn | | |
| 11 | 57505488 | A | C | synonymous_variant | TMX2 | syn | | |
| 11 | 61537815 | A | C | synonymous_variant | MYRF | syn | | |
| 11 | 65403692 | A | G | missense_variant | PCNXL3 | D1836G | 0.077 | 0.051 |
| 11 | 86198475 | A | G | missense_variant | ME3 | L238P | 0.038 | 0.006 |
| 11 | 117078702 | A | C | synonymous_variant | PCSK7 | syn | | |
| 12 | 11420895 | C | T | synonymous_variant | PRB3 | syn | | |
| 12 | 11420962 | C | G | missense_variant | PRB3 | R74P | 0.038 | 0.01 |
| 12 | 11461675 | C | G | missense_variant | PRB4 | R81P | 0.017 | 0.011 |
| 12 | 11506412 | A | T | intron_variant | PRB1 | -- | 0.002 | 0.004 |
| 12 | 11546184 | A | G | synonymous_variant | PRB2 | syn | | |
| 12 | 21327626 | A | C | synonymous_variant | SLCO1B1 | syn | | |
| 12 | 50747002 | C | T | missense_variant | FAM186A | V1205I | 0.039 | 0.01 |
| 12 | 50747044 | A | G | synonymous_variant | FAM186A | syn | | |
| 12 | 50747250 | A | G | missense_variant | FAM186A | L1122P | 0.061 | 0.014 |
| 12 | 50747256 | T | C | missense_variant | FAM186A | E1120G | 0.042 | 0.01 |
| 12 | 53342968 | C | T | missense_variant | KRT18 | T4I | 0.106 | 0.041 |
| 12 | 53343005 | G | A | synonymous_variant | KRT18 | syn | | |
| 12 | 53343059 | C | A | missense_variant | KRT18 | S34R | 0.162 | 0.067 |
| 12 | 53343069 | G | T | missense_variant | KRT18 | G38C | 0.201 | 0.081 |
| 12 | 53343071 | C | T | synonymous_variant | KRT18 | syn | | |
| 12 | 53343074 | T | C | synonymous_variant | KRT18 | syn | | |
| 12 | 132445261 | A | C | missense_variant | EP400 | N33H | 0.112 | 0.138 |
| 14 | 23858236 | T | G | missense_variant | MYH6 | Q1336P | 0.086 | 0.062 |
| 14 | 51348302 | C | T | intron_variant | ABHD12B | -- | | |
| 15 | 20740111 | A | T | missense_variant | GOLGA6L6 | W547R | 0.014 | 0.005 |
| 15 | 42371752 | A | T | missense_variant | PLA2G4D | S434T | 0.012 | 0.008 |
| 15 | 42439376 | G | A | synonymous_variant | PLA2G4F | syn | | |
| 15 | 59515310 | T | C | synonymous_variant | MYO1E | syn | | |
| 15 | 89417238 | A | G | missense_variant | ACAN | Q2500R | 0.034 | 0.074 |
| 15 | 90320185 | A | G | synonymous_variant | MESP2 | syn | | |
| 15 | 90628535 | A | G | missense_variant | IDH2 | V351A | 0.365 | 0.062 |
| 15 | 90628543 | A | C | missense_variant | IDH2 | H348Q | 0.449 | 0.076 |
| 15 | 91432604 | G | A | missense_variant | FES | R246Q | 0.104 | 0.041 |
| 15 | 98504326 | A | G | missense_variant | ARRDC4 | T79A | 0.007 | 0.007 |
| 16 | 2121833 | T | C | synonymous_variant | TSC2 | syn | | |
| 16 | 2121842 | G | C | synonymous_variant | TSC2 | syn | | |
| 16 | 2121860 | T | G | synonymous_variant | TSC2 | syn | | |
| 16 | 29676355 | A | C | 3_prime_UTR_variant | SPN | -- | | |
| 16 | 30016636 | T | C | missense_variant | INO80E | L203P | 0.093 | 0.034 |
| 16 | 30990561 | T | C | missense_variant | SETD1A | S1152P | 0.133 | 0.056 |
| 16 | 85105391 | A | G | missense_variant,splice_region_variant | KIAA0513 | E111G | 0.045 | 0.007 |
| 16 | 89940254 | A | C | missense_variant | TCF25 | N60T | 0.035 | 0.007 |
| 17 | 1373620 | C | G | missense_variant | MYO1C | R792P | 0.088 | 0.049 |
| 17 | 30816606 | G | T | 3_prime_UTR_variant | CDK5R1 | -- | | |
| 17 | 34171597 | A | G | missense_variant | TAF15 | S432G | 0.154 | 0.042 |
| 17 | 39240627 | T | C | missense_variant | KRTAP4-7 | S57P | 0.01 | 0.03 |
| 17 | 39254061 | A | G | synonymous_variant | KRTAP4-8 | syn | | |
| 17 | 39280162 | A | G | synonymous_variant | KRTAP4-12 | syn | | |
| 17 | 45219739 | T | C | missense_variant | CDC27 | K418E | 0.079 | 0.068 |
| 17 | 45219746 | T | G | synonymous_variant | CDC27 | syn | | |
| 17 | 45219750 | T | C | missense_variant | CDC27 | K414R | 0.101 | 0.087 |
| 17 | 45219792 | T | C | missense_variant | CDC27 | K400R | 0.086 | 0.074 |
| 17 | 45219794 | G | C | missense_variant | CDC27 | S399R | 0.096 | 0.083 |
| 17 | 45232068 | A | G | synonymous_variant | CDC27 | syn | | |
| 17 | 61904913 | C | G | 5_prime_UTR_variant | FTSJ3 | -- | | |
| 17 | 72340973 | G | A | missense_variant | KIF19 | R219H | 0.037 | 0.043 |
| 18 | 44140065 | A | C | upstream_gene_variant | LOXHD1 | -- | | |
| 19 | 1881453 | A | C | missense_variant | ABHD17A | V38G | 0.026 | 0.018 |
| 19 | 14272183 | T | G | missense_variant | LPHN1 | Q489P | 0.033 | 0.022 |
| 19 | 39330841 | G | C | synonymous_variant | HNRNPL | syn | | |
| 19 | 47575736 | T | G | missense_variant | ZC3H4 | M559L | 0.082 | 0.137 |
| 19 | 48305612 | T | A | missense_variant | TPRX1 | N219I | 0.014 | 0.005 |
| 19 | 48305634 | A | G | missense_variant | TPRX1 | S212P | 0.011 | 0.004 |
| 19 | 49513773 | A | G | missense_variant | RUVBL2 | E231G | 0.225 | 0.147 |
| 19 | 52887603 | G | A | missense_variant | ZNF880 | R257Q | 0.007 | 0.001 |
| 19 | 54746382 | C | T | missense_variant | LILRA6 | R20H | | |

| | | | | | | | | |
|---|---|---|---|---|---|---|---|---|
| 19 | 54746417 | G | A | intron variant | LILRA6 | -- | | |
| 19 | 58385748 | G | A | missense variant | ZNF814 | A337V | 0.011 | 0.011 |
| 19 | 58385762 | C | G | synonymous variant | ZNF814 | syn | | |
| 20 | 279793 | A | G | 3 prime UTR variant | ZCCHC3 | -- | | |
| 20 | 4708790 | A | C | 3 prime UTR variant | PRND | -- | | |
| 20 | 5548204 | T | A | missense variant,splice region variant | GPCPD1 | K384N | 0.028 | 0.011 |
| 20 | 17640082 | A | G | intron variant | RRBP1 | -- | | |
| 20 | 44512736 | T | C | 3 prime UTR variant | ZSWIM1 | -- | | |
| 20 | 52645325 | T | C | missense variant | BCAS1 | Q110R | 0.004 | 0.008 |
| 20 | 57266792 | G | A | upstream gene variant | NPEPL1 | -- | | |
| 20 | 60982118 | G | C | missense variant | CABLES2 | P72R | 0.055 | 0.014 |
| 20 | 61444665 | G | A | synonymous variant | OGFR | syn | | |
| 20 | 61452837 | G | A | intron variant | COL9A3 | -- | | |
| 20 | 61512376 | G | A | synonymous variant | DIDO1 | syn | | |
| 20 | 62126185 | A | G | synonymous variant | EEF1A2 | syn | | |
| 20 | 62195220 | C | T | missense variant | HELZ2 | R1652Q | 0.021 | 0.019 |
| 20 | 62421622 | A | G | synonymous variant | ZBTB46 | syn | | |
| 21 | 42551245 | A | G | intron variant | BACE2 | -- | | |
| 21 | 42551356 | T | G | intron variant | BACE2 | -- | | |
| 21 | 43531320 | T | C | missense variant | UMODL1 | L663P | 0.038 | 0.024 |
| 21 | 45994187 | C | T | synonymous variant | KRTAP10-4 | syn | | |
| 22 | 21570762 | A | G | missense variant,splice region variant | GGT2 | Y193H | 0.006 | 0.029 |
| 22 | 29885567 | A | C | synonymous variant | NEFH | syn | | |
| 22 | 39777865 | G | T | missense variant | SYNGR1 | Q216H | 0.008 | 0.02 |
| X | 3228653 | G | C | missense variant | MXRA5 | L2531V | 0.015 | 0.021 |
| X | 8434256 | A | G | synonymous variant | VCX3B | syn | | |
| X | 53108600 | T | G | 3 prime UTR variant | GPR173 | -- | | |
| X | 53108621 | T | G | 3 prime UTR variant | GPR173 | -- | | |
| X | 123654613 | T | A | missense variant | TENM1 | I1019F | 0.027 | 0.032 |
| X | 132888189 | C | A | missense variant | GPC3 | V118F | 0.043 | 0.114 |

**[B]** Somatic mutations exclusive to tumorigenic cells were identified by whole exome sequencing. Mutations with variant allele frequencies (VAFs) less than 5% were removed. CHASMplus scores were calculated based on all 32 cancer types and on breast cancer alone.

| Chr | Position | Reference Allele | Alternate Allele | Mutation Consequence | Gene symbol | Amino Acid change | CHASMplus | CHASMplus_BRCA |
|---|---|---|---|---|---|---|---|---|
| 1 | 1132966 | C | A | synonymous variant | TTLL10 | syn | | |
| 1 | 11133978 | C | A | intron variant | EXOSC10 | -- | | |
| 1 | 16388642 | G | A | missense variant | FAM131C | R74C | 0.072 | 0.016 |
| 1 | 21945530 | C | T | synonymous variant | RAP1GAP | syn | | |
| 1 | 86965419 | T | C | synonymous variant | CLCA1 | syn | | |
| 1 | 118166783 | C | T | 3 prime UTR variant | FAM46C | -- | | |
| 1 | 152760431 | G | T | 3 prime UTR variant | LCE1E | -- | | |
| 1 | 243663066 | C | G | missense variant | SDCCAG8 | S707R | 0.005 | 0.002 |
| 2 | 1099077 | A | C | intron variant | SNTG2 | -- | | |
| 2 | 47641557 | G | A | splice region variant,synonymous variant | MSH2 | syn | | |
| 2 | 95542340 | A | G | synonymous variant | TEKT4 | syn | | |
| 2 | 95542344 | C | T | stop gained | TEKT4 | R380* | | |
| 2 | 122205003 | C | A | missense variant | CLASP1 | A608S | 0.076 | 0.057 |
| 2 | 210837831 | G | T | missense variant | UNC80 | Q2742H | 0.052 | 0.074 |
| 2 | 224640645 | C | A | synonymous variant | AP1S3 | syn | | |
| 3 | 10128927 | G | T | missense variant | FANCD2 | A1149S | 0.1 | 0.205 |
| 3 | 77526646 | C | A | missense variant | ROBO2 | P173H | 0.054 | 0.056 |
| 3 | 152881392 | T | G | 3 prime UTR variant | RAP2B | -- | | |
| 4 | 17812796 | C | T | synonymous variant | NCAPG | syn | | |
| 4 | 90874252 | G | T | missense variant | MMRN1 | V1124F | 0.02 | 0.009 |
| 4 | 100137333 | C | T | synonymous variant | ADH6 | syn | | |
| 5 | 31405854 | C | A | intron variant | DROSHA | -- | | |
| 5 | 75427623 | G | A | synonymous variant | SV2C | syn | | |
| 5 | 96101029 | G | A | splice region variant,synonymous variant | CAST | syn | | |
| 5 | 113798802 | C | G | stop gained | KCNN2 | S353* | | |
| 5 | 137088931 | G | A | synonymous variant | HNRNPA0 | syn | | |
| 5 | 140237041 | G | T | missense variant | PCDHA10 | G470C | 0.02 | 0.011 |
| 5 | 172578575 | C | T | missense variant | BNIP1 | L62F | 0.022 | 0.02 |
| 6 | 31648115 | A | C | missense variant | LY6G5C | S12R | 0.021 | 0.021 |
| 6 | 33377484 | G | C | 3 prime UTR variant | KIFC1 | -- | | |
| 6 | 116427489 | G | T | missense variant | NT5DC1 | W59L | 0.081 | 0.011 |
| 6 | 154762437 | T | A | missense variant | CNKSR3 | T166S | 0.009 | 0.019 |
| 7 | 77006706 | A | C | missense variant,splice region variant | GSAP | V193G | 0.045 | 0.03 |
| 7 | 83277744 | C | T | missense variant,splice region variant | SEMA3E | E39K | 0.004 | 0.006 |
| 9 | 21351376 | C | A | 5 prime UTR variant | IFNA6 | -- | | |
| 9 | 32986010 | T | C | missense variant | APTX | S168G | 0.011 | 0.015 |
| 9 | 96422719 | T | A | missense variant | PHF2 | D525E | 0.016 | 0.023 |
| 9 | 111870734 | T | A | missense variant,splice region variant | TMEM245 | L232F | 0.019 | 0.015 |
| 9 | 111870738 | T | A | missense variant | TMEM245 | H231L | 0.037 | 0.03 |
| 10 | 43610096 | T | A | missense variant | RET | F683Y | 0.075 | 0.04 |

| | | | | | | | | |
|---|---|---|---|---|---|---|---|---|
| 10 | 101295196 | C | G | synonymous_variant | NKX2-3 | syn | | |
| 10 | 115805564 | T | C | 3_prime_UTR_variant | ADRB1 | -- | | |
| 11 | 1651127 | C | T | synonymous_variant | KRTAP5-5 | syn | | |
| 11 | 44286566 | G | A | synonymous_variant | ALX4 | syn | | |
| 11 | 46342327 | C | A | 3_prime_UTR_variant | CREB3L1 | -- | | |
| 11 | 57311355 | A | G | synonymous_variant | SMTNL1 | syn | | |
| 11 | 64074764 | C | T | missense_variant | ESRRA | A38V | 0.063 | 0.038 |
| 11 | 102822868 | G | T | missense_variant | MMP13 | F224L | 0.025 | 0.004 |
| 11 | 108014696 | C | T | intron_variant | ACAT1 | -- | | |
| 11 | 117097952 | C | T | synonymous_variant | PCSK7 | syn | | |
| 11 | 118255660 | G | A | splice_region_variant,synonymous_variant | UBE4A | syn | | |
| 11 | 123676980 | G | T | missense_variant | OR6M1 | F26L | 0.003 | 0.008 |
| 12 | 48174336 | C | T | 3_prime_UTR_variant | SLC48A1 | -- | | |
| 12 | 57642890 | G | A | missense_variant | STAC3 | H90Y | 0.073 | 0.09 |
| 12 | 76447645 | T | A | synonymous_variant | NAP1L1 | syn | | |
| 12 | 129189941 | G | A | missense_variant | TMEM132C | G810R | 0.013 | 0.011 |
| 12 | 132547078 | G | A | synonymous_variant | EP400 | syn | | |
| 12 | 132547081 | G | A | synonymous_variant | EP400 | syn | | |
| 13 | 20077348 | T | C | intron_variant | TPTE2 | -- | | |
| 13 | 20077352 | T | C | intron_variant | TPTE2 | -- | | |
| 13 | 51287408 | C | G | intron_variant | DLEU7 | -- | | |
| 13 | 98658582 | C | T | synonymous_variant | IPO5 | syn | | |
| 14 | 21854004 | T | G | missense_variant | CHD8 | H2505P | 0.398 | 0.113 |
| 15 | 34659253 | C | G | missense_variant | LPCAT4 | G17R | 0.046 | 0.072 |
| 15 | 80216931 | T | C | 3_prime_UTR_variant | C15orf37 | -- | | |
| 16 | 3639856 | C | T | synonymous_variant | SLX4 | syn | | |
| 16 | 23722280 | G | A | synonymous_variant | ERN2 | syn | | |
| 16 | 27752135 | C | T | synonymous_variant | KIAA0556 | syn | | |
| 17 | 10432122 | C | T | missense_variant | MYH2 | G1210E | 0.066 | 0.033 |
| 17 | 25638634 | A | T | missense_variant | WSB1 | L366F | 0.028 | 0.019 |
| 17 | 37863247 | C | T | synonymous_variant | ERBB2 | syn | | |
| 17 | 37866418 | T | C | synonymous_variant | ERBB2 | syn | | |
| 17 | 37866421 | C | A | synonymous_variant | ERBB2 | syn | | |
| 17 | 37866442 | C | T | synonymous_variant | ERBB2 | syn | | |
| 17 | 37868601 | T | C | synonymous_variant | ERBB2 | syn | | |
| 17 | 37868603 | G | T | missense_variant | ERBB2 | L350F | 0.211 | 0.192 |
| 17 | 37868608 | A | G | missense_variant | ERBB2 | E352G | 0.115 | 0.141 |
| 17 | 37868611 | T | C | missense_variant | ERBB2 | V353A | 0.164 | 0.141 |
| 17 | 37868618 | A | C | synonymous_variant | ERBB2 | syn | | |
| 17 | 37868619 | G | A | missense_variant | ERBB2 | V356I | 0.175 | 0.192 |
| 17 | 37868621 | T | C | synonymous_variant | ERBB2 | syn | | |
| 17 | 37868629 | C | A | missense_variant | ERBB2 | A359D | 0.199 | 0.204 |
| 17 | 37868634 | A | G | missense_variant | ERBB2 | I361V | 0.206 | 0.202 |
| 17 | 37868647 | C | A | missense_variant | ERBB2 | A365D | 0.19 | 0.199 |
| 17 | 37868682 | C | T | synonymous_variant | ERBB2 | syn | | |
| 17 | 37879808 | A | G | synonymous_variant | ERBB2 | syn | | |
| 17 | 37879811 | T | C | synonymous_variant | ERBB2 | syn | | |
| 17 | 37879820 | G | A | synonymous_variant | ERBB2 | syn | | |
| 17 | 37879835 | G | T | synonymous_variant | ERBB2 | syn | | |
| 17 | 37879850 | G | A | synonymous_variant | ERBB2 | syn | | |
| 17 | 37879865 | G | A | synonymous_variant | ERBB2 | syn | | |
| 17 | 37879889 | T | A | synonymous_variant | ERBB2 | syn | | |
| 17 | 37879892 | C | A | synonymous_variant | ERBB2 | syn | | |
| 17 | 37879904 | A | T | synonymous_variant | ERBB2 | syn | | |
| 17 | 37880179 | T | A | synonymous_variant | ERBB2 | syn | | |
| 17 | 37880200 | T | C | synonymous_variant | ERBB2 | syn | | |
| 17 | 37880203 | A | C | synonymous_variant | ERBB2 | syn | | |
| 17 | 37880209 | C | T | synonymous_variant | ERBB2 | syn | | |
| 17 | 37880215 | A | G | synonymous_variant | ERBB2 | syn | | |
| 17 | 37880224 | G | A | synonymous_variant | ERBB2 | syn | | |
| 17 | 37880236 | C | T | synonymous_variant | ERBB2 | syn | | |
| 17 | 37880239 | C | T | synonymous_variant | ERBB2 | syn | | |
| 17 | 37881005 | C | T | synonymous_variant | ERBB2 | syn | | |
| 17 | 37881008 | C | T | synonymous_variant | ERBB2 | syn | | |
| 17 | 37881011 | A | G | synonymous_variant | ERBB2 | syn | | |
| 17 | 37881017 | C | G | synonymous_variant | ERBB2 | syn | | |
| 17 | 37881026 | T | C | synonymous_variant | ERBB2 | syn | | |
| 17 | 37881050 | G | A | synonymous_variant | ERBB2 | syn | | |
| 17 | 37881053 | G | A | synonymous_variant | ERBB2 | syn | | |
| 17 | 37881080 | T | C | synonymous_variant | ERBB2 | syn | | |
| 17 | 37881089 | C | T | synonymous_variant | ERBB2 | syn | | |
| 17 | 37881090 | T | C | synonymous_variant | ERBB2 | syn | | |
| 17 | 37881092 | A | G | synonymous_variant | ERBB2 | syn | | |
| 17 | 37881104 | G | A | synonymous_variant | ERBB2 | syn | | |
| 17 | 37881108 | A | C | missense_variant | ERBB2 | N813H | 0.482 | 0.42 |
| 17 | 37881113 | C | A | synonymous_variant | ERBB2 | syn | | |

| | | | | | | | | |
|---|---|---|---|---|---|---|---|---|
| 17 | 37881116 | A | T | synonymous_variant | ERBB2 | syn | | |
| 17 | 37881122 | G | A | synonymous_variant | ERBB2 | syn | | |
| 17 | 37881140 | G | C | synonymous_variant | ERBB2 | syn | | |
| 17 | 37881150 | A | G | missense_variant | ERBB2 | M827V | 0.255 | 0.28 |
| 17 | 37881152 | G | T | missense_variant | ERBB2 | M827I | 0.258 | 0.288 |
| 17 | 37881322 | T | C | synonymous_variant | ERBB2 | syn | | |
| 17 | 37881331 | C | T | synonymous_variant | ERBB2 | syn | | |
| 17 | 37881344 | T | C | synonymous_variant | ERBB2 | syn | | |
| 17 | 37881349 | C | T | synonymous_variant | ERBB2 | syn | | |
| 17 | 37881352 | T | C | synonymous_variant | ERBB2 | syn | | |
| 17 | 37881358 | C | T | synonymous_variant | ERBB2 | syn | | |
| 17 | 37881364 | G | A | synonymous_variant | ERBB2 | syn | | |
| 17 | 37881382 | T | C | synonymous_variant | ERBB2 | syn | | |
| 17 | 37881388 | A | G | synonymous_variant | ERBB2 | syn | | |
| 17 | 37881397 | C | T | synonymous_variant | ERBB2 | syn | | |
| 17 | 37881427 | C | T | synonymous_variant | ERBB2 | syn | | |
| 17 | 37881600 | G | A | synonymous_variant | ERBB2 | syn | | |
| 17 | 37881633 | C | T | synonymous_variant | ERBB2 | syn | | |
| 17 | 37881651 | T | C | synonymous_variant | ERBB2 | syn | | |
| 17 | 37882009 | G | A | synonymous_variant | ERBB2 | syn | | |
| 17 | 37882033 | C | T | synonymous_variant | ERBB2 | syn | | |
| 17 | 37882034 | C | T | synonymous_variant | ERBB2 | syn | | |
| 17 | 37882042 | A | G | synonymous_variant | ERBB2 | syn | | |
| 17 | 37882048 | G | A | synonymous_variant | ERBB2 | syn | | |
| 17 | 37882051 | G | A | synonymous_variant | ERBB2 | syn | | |
| 17 | 37882054 | G | C | synonymous_variant | ERBB2 | syn | | |
| 17 | 37882057 | G | A | synonymous_variant | ERBB2 | syn | | |
| 17 | 37882060 | C | T | synonymous_variant | ERBB2 | syn | | |
| 17 | 37882066 | C | T | synonymous_variant | ERBB2 | syn | | |
| 17 | 37882069 | C | A | synonymous_variant | ERBB2 | syn | | |
| 17 | 37882096 | C | T | synonymous_variant | ERBB2 | syn | | |
| 17 | 37882840 | G | C | synonymous_variant | ERBB2 | syn | | |
| 17 | 37882843 | A | G | synonymous_variant | ERBB2 | syn | | |
| 17 | 37882864 | T | A | synonymous_variant | ERBB2 | syn | | |
| 17 | 37882870 | C | T | synonymous_variant | ERBB2 | syn | | |
| 17 | 37882876 | C | T | synonymous_variant | ERBB2 | syn | | |
| 17 | 37882882 | C | G | synonymous_variant | ERBB2 | syn | | |
| 17 | 37882897 | C | T | synonymous_variant | ERBB2 | syn | | |
| 17 | 37883070 | T | C | splice_region_variant,synonymous_variant | ERBB2 | syn | | |
| 17 | 37883086 | G | T | missense_variant | ERBB2 | A997S | 0.26 | 0.445 |
| 17 | 37883091 | T | C | synonymous_variant | ERBB2 | syn | | |
| 17 | 37883095 | T | A | missense_variant | ERBB2 | L1000M | 0.208 | 0.204 |
| 17 | 37883103 | C | T | synonymous_variant | ERBB2 | syn | | |
| 17 | 37883115 | C | T | synonymous_variant | ERBB2 | syn | | |
| 17 | 37883127 | G | A | synonymous_variant | ERBB2 | syn | | |
| 17 | 37883130 | C | T | synonymous_variant | ERBB2 | syn | | |
| 17 | 37883142 | G | T | synonymous_variant | ERBB2 | syn | | |
| 17 | 37883151 | G | A | synonymous_variant | ERBB2 | syn | | |
| 17 | 37883154 | T | C | synonymous_variant | ERBB2 | syn | | |
| 17 | 37883160 | G | A | synonymous_variant | ERBB2 | syn | | |
| 17 | 37883172 | A | G | synonymous_variant | ERBB2 | syn | | |
| 17 | 37883553 | C | A | synonymous_variant | ERBB2 | syn | | |
| 17 | 37883559 | G | T | synonymous_variant | ERBB2 | syn | | |
| 17 | 37883562 | C | G | missense_variant | ERBB2 | D1058E | 0.197 | 0.167 |
| 17 | 37883571 | A | G | synonymous_variant | ERBB2 | syn | | |
| 17 | 37883574 | G | C | synonymous_variant | ERBB2 | syn | | |
| 17 | 37883586 | T | G | synonymous_variant | ERBB2 | syn | | |
| 17 | 37883592 | G | A | synonymous_variant | ERBB2 | syn | | |
| 17 | 37883594 | A | G | missense_variant | ERBB2 | E1069G | 0.185 | 0.16 |
| 17 | 37883596 | G | C | missense_variant | ERBB2 | A1070P | 0.217 | 0.211 |
| 17 | 37883604 | G | A | synonymous_variant | ERBB2 | syn | | |
| 17 | 37883616 | A | T | synonymous_variant | ERBB2 | syn | | |
| 17 | 37883622 | C | G | synonymous_variant | ERBB2 | syn | | |
| 17 | 37883643 | A | G | synonymous_variant | ERBB2 | syn | | |
| 17 | 37883660 | G | C | missense_variant | ERBB2 | G1091A | 0.166 | 0.16 |
| 17 | 37883669 | C | T | missense_variant | ERBB2 | A1094V | 0.131 | 0.121 |
| 17 | 37883671 | G | A | missense_variant | ERBB2 | A1095T | 0.145 | 0.13 |
| 17 | 37883676 | G | A | synonymous_variant | ERBB2 | syn | | |
| 17 | 37883685 | A | G | synonymous_variant | ERBB2 | syn | | |
| 17 | 37883692 | C | T | missense_variant | ERBB2 | P1102S | 0.171 | 0.155 |
| 17 | 37883694 | C | T | synonymous_variant | ERBB2 | syn | | |
| 17 | 37883695 | A | C | missense_variant | ERBB2 | T1103P | 0.229 | 0.206 |
| 17 | 37883705 | C | T | missense_variant | ERBB2 | P1106L | 0.178 | 0.181 |
| 17 | 37883727 | T | C | synonymous_variant | ERBB2 | syn | | |
| 17 | 37883740 | G | T | missense_variant | ERBB2 | V1118L | 0.23 | 0.218 |
| 17 | 37883745 | C | T | synonymous_variant | ERBB2 | syn | | |

| | | | | | | | | |
|---|---|---|---|---|---|---|---|---|
| 17 | 37883754 | T | C | synonymous variant | ERBB2 | syn | | |
| 17 | 37883769 | C | T | synonymous variant | ERBB2 | syn | | |
| 17 | 37883775 | C | T | synonymous variant | ERBB2 | syn | | |
| 17 | 37883782 | A | G | missense variant | ERBB2 | T1132A | 0.134 | 0.12 |
| 17 | 37883994 | C | T | synonymous variant | ERBB2 | syn | | |
| 17 | 37884004 | G | C | missense variant | ERBB2 | A1159P | 0.175 | 0.186 |
| 17 | 37884008 | C | T | missense variant | ERBB2 | A1160V | 0.112 | 0.121 |
| 17 | 37884012 | A | G | synonymous variant | ERBB2 | syn | | |
| 17 | 37884024 | C | T | synonymous variant | ERBB2 | syn | | |
| 17 | 37884030 | G | A | synonymous variant | ERBB2 | syn | | |
| 17 | 37884036 | G | A | synonymous variant | ERBB2 | syn | | |
| 17 | 37884051 | C | T | synonymous variant | ERBB2 | syn | | |
| 17 | 37884054 | A | T | synonymous variant | ERBB2 | syn | | |
| 17 | 37884069 | C | T | synonymous variant | ERBB2 | syn | | |
| 17 | 37884090 | T | C | synonymous variant | ERBB2 | syn | | |
| 17 | 37884099 | C | T | synonymous variant | ERBB2 | syn | | |
| 17 | 37884111 | C | T | synonymous variant | ERBB2 | syn | | |
| 17 | 37884114 | G | A | synonymous variant | ERBB2 | syn | | |
| 17 | 37884146 | A | C | missense variant | ERBB2 | Q1206P | 0.258 | 0.234 |
| 17 | 37884157 | C | T | missense variant | ERBB2 | P1210S | 0.218 | 0.213 |
| 17 | 37884180 | C | T | synonymous variant | ERBB2 | syn | | |
| 17 | 37884205 | G | A | missense variant | ERBB2 | D1226N | 0.164 | 0.167 |
| 17 | 37884208 | C | T | missense variant | ERBB2 | P1227S | 0.19 | 0.144 |
| 17 | 44851038 | G | T | missense variant | WNT3 | D106E | 0.016 | 0.015 |
| 19 | 7584223 | A | G | missense variant | ZNF358 | N32S | 0.025 | 0.005 |
| 19 | 9006729 | G | A | synonymous variant | MUC16 | syn | | |
| 19 | 9024951 | T | A | missense variant | MUC16 | H12304L | 0.013 | 0.009 |
| 19 | 30106173 | G | T | synonymous variant | POP4 | syn | | |
| 19 | 38993547 | C | T | synonymous variant | RYR1 | syn | | |
| 20 | 32664866 | G | A | missense variant | RALY | G231S | 0.049 | 0.058 |
| 20 | 33033099 | C | G | missense variant | ITCH | R366G | 0.321 | 0.226 |
| 21 | 42551433 | G | A | intron variant | BACE2 | -- | | |
| 22 | 37329999 | C | T | synonymous variant | CSF2RB | syn | | |
| 22 | 42276786 | A | T | missense variant | SREBF2 | T610S | 0.169 | 0.096 |
| X | 49454131 | G | C | missense variant | PAGE1 | A103G | 0.004 | 0.002 |
| X | 102864007 | C | A | stop gained | TCEAL3 | Y5* | | |
| X | 149887114 | T | A | missense variant | MTMR1 | N90K | 0.078 | 0.028 |
| X | 153048675 | T | C | splice region variant,intron variant | SRPK3 | -- | | |

**[C]** Among somatic mutations that are shared between tumorigenic and non-tumorigenic cells, variant allele frequencies (VAFs) that are greater by at least 1.5-fold in tumorigenic cells than in non-tumorigenic cells are presented. The mutations were identified by whole exome sequencing. Mutations with variant allele frequencies less than 5% were removed. CHASMplus scores were calculated based on all 32 cancer types and on breast cancer alone.

| Chr | Position | Reference Allele | Alternate Allele | Mutation Consequence | Gene symbol | Amino Acid change | CHASMplus | CHASMplus_BRCA | Fold-change (tumor/non-tumor) |
|---|---|---|---|---|---|---|---|---|---|
| 1 | 67591575 | A | C | missense variant | C1orf141 | S31R | 0.001 | 0.004 | 1.642 |
| 1 | 144852518 | C | T | intron variant | PDE4DIP | -- | | | 1.730 |
| 1 | 144852522 | C | T | intron variant | PDE4DIP | -- | | | 1.548 |
| 1 | 156873727 | G | A | synonymous variant | PEAR1 | syn | | | 2.066 |
| 1 | 200880978 | C | T | missense variant | C1orf106 | R453C | 0.009 | 0.011 | 1.767 |
| 1 | 224577599 | C | G | intron variant | WDR26 | -- | | | 1.625 |
| 1 | 243736228 | C | T | splice region variant,synonymous variant | AKT3 | syn | | | 1.534 |
| 2 | 95542376 | G | T | synonymous variant | TEKT4 | syn | | | 2.850 |
| 2 | 95542377 | C | T | missense variant | TEKT4 | R391C | 0.007 | 0.008 | 2.850 |
| 2 | 95542419 | G | A | missense variant | TEKT4 | A405T | 0.002 | 0.004 | 2.000 |
| 2 | 231036866 | G | A | synonymous variant | SP110 | syn | | | 1.938 |
| 2 | 240982164 | T | C | missense variant | PRR21 | H79R | | | 1.641 |
| 3 | 152881416 | T | G | 3 prime UTR variant | RAP2B | -- | | | 1.817 |
| 3 | 160253665 | G | A | synonymous variant | KPNA4 | syn | | | 1.653 |
| 3 | 195513946 | A | G | missense variant | MUC4 | V1502A | 0.036 | 0.022 | 1.600 |
| 4 | 57180616 | G | C | synonymous variant | KIAA1211 | syn | | | 1.662 |
| 5 | 98216991 | C | T | missense variant | CHD1 | E986K | 0.164 | 0.068 | 1.504 |
| 5 | 139496402 | G | T | downstream gene variant | PURA | -- | | | 2.857 |
| 5 | 175957139 | C | G | synonymous variant | RNF44 | syn | | | 1.630 |
| 6 | 21597780 | G | T | 3 prime UTR variant | SOX4 | -- | | | 1.931 |
| 6 | 21597800 | A | T | 3 prime UTR variant | SOX4 | -- | | | 1.568 |
| 6 | 30954414 | C | G | missense variant | MUC21 | D154E | 0.025 | 0.018 | 1.531 |
| 6 | 30954684 | G | C | missense variant | MUC21 | E244D | 0.027 | 0.018 | 2.636 |
| 6 | 30954694 | C | A | missense variant | MUC21 | P248T | 0.024 | 0.015 | 2.807 |
| 6 | 30954729 | G | C | missense variant | MUC21 | E259D | 0.027 | 0.018 | 1.756 |
| 6 | 30954749 | G | A | missense variant | MUC21 | G266E | 0.028 | 0.018 | 1.504 |
| 6 | 30954939 | A | T | synonymous variant | MUC21 | syn | | | 3.400 |
| 6 | 33167066 | T | G | synonymous variant | RXRB | syn | | | 1.528 |

| | | | | | | | | | |
|---|---|---|---|---|---|---|---|---|---|
| 6 | 122772903 | C | A | splice_region_variant,intron_variant | SERINC1 | -- | | | 1.558 |
| 6 | 160679400 | C | A | synonymous_variant | SLC22A2 | syn | | | 1.586 |
| 6 | 167595371 | T | C | synonymous_variant | TCP10L2 | syn | | | 2.857 |
| 7 | 75597545 | C | T | intron_variant | POR | -- | | | 1.851 |
| 7 | 77006694 | G | A | missense_variant | GSAP | S197F | 0.038 | 0.026 | 4.125 |
| 7 | 86556221 | C | A | missense_variant | KIAA1324L | M367I | 0.018 | 0.013 | 1.587 |
| 7 | 103202398 | G | A | missense_variant,splice_region_variant | RELN | S1738F | 0.054 | 0.029 | 1.598 |
| 7 | 143956461 | T | C | synonymous_variant | OR2A7 | syn | | | 1.743 |
| 8 | 142228149 | G | A | synonymous_variant | SLC45A4 | syn | | | 1.505 |
| 9 | 69247530 | T | A | missense_variant | CBWD6 | Y161F | | | 3.318 |
| 9 | 131284948 | A | G | missense_variant,splice_region_variant | GLE1 | E145G | 0.056 | 0.026 | 1.852 |
| 9 | 139008644 | G | A | missense_variant | C9orf69 | R35C | 0.014 | 0.019 | 2.011 |
| 10 | 93489 | G | C | stop_gained | TUBB8 | Y281* | | | 1.521 |
| 10 | 5567967 | G | T | 3_prime_UTR_variant | CALML3 | -- | | | 1.962 |
| 10 | 21807832 | A | T | 5_prime_UTR_variant | SKIDA1 | -- | | | 1.631 |
| 10 | 81273535 | T | A | 3_prime_UTR_variant | EIF5AL1 | -- | | | 1.954 |
| 10 | 99504630 | G | T | missense_variant | ZFYVE27 | G138V | 0.085 | 0.038 | 1.857 |
| 11 | 1093286 | C | G | missense_variant | MUC2 | T1702S | | | 2.700 |
| 11 | 1093364 | C | G | missense_variant | MUC2 | T1728S | | | 1.870 |
| 11 | 1093368 | G | A | synonymous_variant | MUC2 | syn | | | 1.762 |
| 11 | 1093575 | G | T | synonymous_variant | MUC2 | syn | | | 2.299 |
| 11 | 64415767 | G | A | synonymous_variant | NRXN2 | syn | | | 1.518 |
| 12 | 11461745 | G | C | missense_variant | PRB4 | Q58E | 0.015 | 0.011 | 1.972 |
| 12 | 11506774 | C | T | missense_variant | PRB1 | R88Q | 0.003 | 0.004 | 2.869 |
| 12 | 11506785 | C | T | synonymous_variant | PRB1 | syn | | | 3.378 |
| 12 | 51740409 | T | G | missense_variant,splice_region_variant | CELA1 | Y5S | 0.014 | 0.009 | 3.612 |
| 13 | 20000613 | A | G | synonymous_variant | TPTE2 | syn | | | 1.676 |
| 14 | 21167576 | A | T | missense_variant | RNASE4 | T16S | 0.007 | 0.014 | 1.547 |
| 14 | 34269775 | C | G | synonymous_variant | NPAS3 | syn | | | 1.678 |
| 15 | 73545732 | T | C | intron_variant | NEO1 | -- | | | 1.580 |
| 16 | 67267851 | A | G | synonymous_variant | FHOD1 | syn | | | 3.810 |
| 16 | 74425314 | G | A | missense_variant | NPIPB15 | R223H | 0.006 | 0.002 | 1.670 |
| 16 | 74425358 | G | A | missense_variant | NPIPB15 | A238T | 0.003 | 0.001 | 2.193 |
| 16 | 74451970 | G | C | missense_variant | CLEC18B | T148S | 0.009 | 0.024 | 1.557 |
| 16 | 88709828 | A | G | missense_variant | CYBA | V174A | 0.124 | 0.034 | 1.700 |
| 16 | 89662990 | T | C | synonymous_variant | CPNE7 | syn | | | 1.690 |
| 17 | 1733399 | A | G | synonymous_variant | RPA1 | syn | | | 1.607 |
| 17 | 39150329 | T | G | synonymous_variant | KRTAP3-3 | syn | | | 1.840 |
| 17 | 39274087 | G | C | missense_variant | KRTAP4-11 | L161V | 0.007 | 0.006 | 1.734 |
| 19 | 1881395 | T | C | synonymous_variant | ABHD17A | syn | | | 3.846 |
| 19 | 1881408 | A | G | missense_variant | ABHD17A | L53S | 0.022 | 0.017 | 1.783 |
| 19 | 1881530 | G | A | synonymous_variant | ABHD17A | syn | | | 3.264 |
| 19 | 9002623 | C | T | missense_variant | MUC16 | R13398H | 0.019 | 0.014 | 1.839 |
| 19 | 9002631 | G | A | synonymous_variant | MUC16 | syn | | | 2.071 |
| 19 | 9002636 | C | T | missense_variant | MUC16 | V13394I | 0.023 | 0.015 | 2.069 |
| 19 | 9002637 | A | G | synonymous_variant | MUC16 | syn | | | 2.107 |
| 19 | 39227921 | G | C | missense_variant | CAPN12 | R413G | 0.017 | 0.01 | 1.750 |
| 19 | 48305566 | C | T | synonymous_variant | TPRX1 | syn | | | 2.159 |
| 19 | 55286665 | T | A | missense_variant | KIR2DL1 | L140Q | 0.029 | 0.009 | 1.829 |
| 20 | 20079300 | T | G | missense_variant,splice_region_variant | C20orf26 | V234G | 0.015 | 0.01 | 2.196 |
| 20 | 57045765 | A | G | missense_variant | APCDD1L | C30R | 0.002 | 0.014 | 2.014 |
| 20 | 57429627 | C | A | missense_variant | GNAS | A436D | 0.129 | 0.073 | 1.943 |
| 20 | 60899196 | C | T | missense_variant | LAMA5 | R1903K | 0.02 | 0.017 | 2.898 |
| 20 | 61040951 | C | T | synonymous_variant | GATA5 | syn | | | 2.030 |
| 21 | 45994014 | C | T | missense_variant | KRTAP10-4 | P127S | 0.019 | 0.005 | 1.586 |
| 21 | 46012181 | C | T | missense_variant | KRTAP10-6 | R62H | 0.017 | 0.014 | 1.632 |
| 21 | 46012182 | G | A | missense_variant | KRTAP10-6 | R62C | 0.02 | 0.017 | 1.650 |
| 21 | 46012345 | G | A | synonymous_variant | KRTAP10-6 | syn | | | 2.129 |
| 21 | 46012351 | G | A | synonymous_variant | KRTAP10-6 | syn | | | 2.642 |
